# A new comparative framework for estimating selection on synonymous substitutions

**DOI:** 10.1101/2024.09.17.613331

**Authors:** Hannah Verdonk, Alyssa Pivirotto, Vitor Pavinato, Jody Hey, Sergei LK Pond

**Affiliations:** Institute for Genomics and Evolutionary Medicine, Department of Biology, Temple University, Philadelphia, Pennsylvania, USA; Center for Computational Genetics and Genomics, Department of Biology, Temple University, Philadelphia, Pennsylvania, USA

## Abstract

Selection on synonymous codon usage is a well known and widespread phenomenon, yet existing models often do not account for it or its effect on synonymous substitution rates. In this article, we develop and expand the capabilities of Multiclass Synonymous Substitution (MSS) models, which account for such selection by partitioning synonymous substitutions into two or more classes and estimating a relative substitution rate for each class, while accounting for important confounders like mutation bias. We identify extensive heterogeneity among relative synonymous substitution rates in an empirical dataset of ∼12,000 gene alignments from twelve *Drosophila* species. We validate model performance using data simulated under a forward population genetic simulation, demonstrating that MSS models are robust to model misspecification. MSS rates are significantly correlated with other covariates of selection on codon usage (population-level polymorphism data and tRNA abundance data), suggesting that models can detect weak signatures of selection on codon usage. With the MSS model, we can now study selection on synonymous substitutions in diverse taxa, independent of any *a priori* assumptions about the forces driving that selection.

## Introduction

A fundamental goal of evolutionary biology is to identify drivers of natural selection acting on genes and genomes. Synonymous or silent substitutions shape codon usage across the genome, and although they were once widely regarded as neutral, the field is gaining an increasing appreciation that such a view is unjustifiably simplistic. Fitness considerations underlying selective forces acting on synonymous changes include translational efficiency (tRNA abundances), successful mRNA folding, and maintenance of CpG sites for methylation (Ikemura 1981; Kanaya et al. 2001; Trotta 2013). Selection on codon usage – a compelling indicator of selection – has been long recognized as widespread in *Drosophila* (Marais et al. 2004), Enterobacteriaceae (P M Sharp and Li 1987), and other taxonomic groups (Marais and Duret 2001; Trotta 2013). Synonymous substitution rates, corrected for nucleotide rate biases and codon frequencies, should not, therefore, be expected to be the same for all codon pairs. This would be conceptually similar to expecting the F81 (Felsenstein 1981) nucleotide substitution model to be biologically adequate. Instead, rates should be allowed to vary among synonymous codon pairs, and be partially informed by key biological traits like gene expression. Such correlations can inform researchers about the relative importance of underlying mechanisms driving selection on synonymous codon usage.

Multiple classic studies conducted in the pre-genomic era found evidence for purifying selection on codon usage as a way to maintain translational efficiency and accuracy in highly expressed genes (Ikemura 1981; Moriyama and Powell 1997; Iriarte et al. 2013). The link between codon usage bias and gene expression is well-studied and supported, but does not (nor should it) fully explain selection on synonymous codon usage. For example, Goetz and Fuglsang found that while gene transcript abundance and various measures of codon usage bias in *E. coli* (Goetz and Fuglsang 2005) were correlated, the largest correlation (with Sharp and Li’s codon adaptation index, CAI (Paul M. Sharp and Li 1987) was only ∼0.43. Clearly, studies that consider how selection for transcriptional/translational efficiency shapes codon usage bias cannot account for other selective pressures on synonymous codon usage. Synonymous substitution rates, corrected for nucleotide rate biases and codon frequencies, should not, therefore, be expected to be the same for all codon pairs. This would be conceptually similar to expecting the F81 (Felsenstein 1981) nucleotide substitution model to be biologically adequate. Instead, rates should be allowed to vary among synonymous codon pairs, and be partially informed by key biological traits like gene expression. Such correlations can inform researchers about the relative importance of underlying mechanisms driving selection on synonymous codon usage.

Kosiol, Holmes, and Goldman took a comparative genomic approach to studying how substitution rates vary among synonymous codon pairs (Kosiol et al. 2007). The authors directly estimated substitution rates for all exchangeable codon pairs from a large dataset of protein-coding gene sequences, and incorporated the numerically estimated rates into an empirical codon evolutionary model. Their model is advantageous in that they directly estimate synonymous substitution rates from input data without assuming an underlying selective pressure, allowing one to explore different biological correlates with substitution rate. However, large training datasets of protein-coding genes may be difficult and time-consuming to obtain for some taxa, and could hinder similar studies in other organisms. The model is also difficult to generalize across genes and across taxa because it aggregates substitution rates across both.

More recently, Rahman *et. al.* explored selection on synonymous substitutions in Enterobacteriaceae (Rahman et al. 2021). In this dataset, Alanine and Valine were the two amino-acids whose codon usage bias was least correlated with expression, and thus assumed to be least likely to be affected by selection on translational efficiency. Defining a Multiple Synonymous Substitution (MSS) codon model which treated synonymous substitutions between A and V codons (both 4-fold degenerate) as neutral, and all others as selected, Rahman *et. al.* were able to show that a weak purifying selection operates on other synonymous substitutions, conditioned on the externally defined set of “neutral” synonymous substitutions. However, it remains unclear how the choice of this neutral set affects the model and inferences, and whether or not all genes are equally agreeable to it.

Here we expand the class of data-driven MSS evolutionary models. An MSS model is a codon-substitution model that includes two or more substitution rate parameters for synonymous codon pairs, in addition to differentiating synonymous and non-synonymous substitutions. Rates are estimated directly from sequence alignments using phylogenetic maximum likelihood. The most complex MSS model, called SynREVCodon, partitions synonymous substitutions into 67 classes - one for each pair of synonymous codons reachable by a single nucleotide substitution, assuming the universal genetic code - and estimates a relative substitution rate for each class. Like other codon models, MSS accounts for important confounders, such as phylogenetic relatedness, nucleotide substitution biases, and unequal codon frequencies. MSS enables a rigorous statistical comparison of synonymous substitution rates within a gene or across a set of genes. We define and apply MSS models of different scope and complexity to a collection of ∼12,000 gene alignments from twelve *Drosophila* species to investigate the extent and correlates of rate variation among synonymous substitutions. We use simulations, population-level data, and tRNA abundance as extrinsic covariates to validate model performance.

## Materials and Methods

### Multiclass Synonymous Substitution (MSS) models

Multiclass Synonymous Substitution (MSS) models extend the standard Muse-Gaut (MG94) class of codon substitution models (Muse and Gaut 1994), by allowing synonymous substitutions to occur at different relative rates. The instantaneous **rate** of substitution from one sense (x) codon to another (y), *x* → *y* is defined as follows.

We follow the standard assumption that multi-nucleotide substitutions must be realized via a sequence of single-step substitutions, although this assumption can be relaxed (Lucaci et al. 2021). θ (*x*, *y*) encodes nucleotide substitutional biases, and follows the MG94xREV convention (Wisotsky et al. 2020), with θ (*x*, *y*) = θ_*nm*_, where **n** and **m** are the two nucleotides being exchanged. Because of time-reversibility, θ (*x*, *y*)= θ (*y*, *x*), and because of identifiability constraints (only products of rates and times can be inferred), we assert θ_*AG*_ = 1. In the Muse-Gaut framework, π (*x*, *y*) is the estimated equilibrium frequency of the target nucleotide of the substitution; we use the corrected empirical F3×4 (CF3×4) estimator (Pond et al. 2010). MSS models differ from standard codon-substitution models in that they allow synonymous substitution rates, α (*x*, *y*), to depend on which codons are being exchanged. We consider several parameterizations of this function and the corresponding models.

For the *a priori* models, one of the substitution rates or groups of rates is designated beforehand as the reference/neutral rate (α_0_) and constrained to 1, both to assist with interpretation and to ensure identifiability. For SynREV and SynREVCodon, the mean of all α (*x*, *y*) is constrained to 1. In this paper, the non-synonymous rate component is parameterized as β (*x*, *y*) : = β, where β is the shared inferred non-synonymous substitution rate that does not depend on the codons. Because α (*x*, *y*) is constrained to have mean 1, β can be interpreted as the standard dN/dS (or ω) ratio. In addition to gene-wide model parameters, a separate branch length parameter is included for every branch in the phylogenetic tree. Unless stated otherwise, all model parameters are estimated by maximum likelihood, using direct optimization, as implemented in the HyPhy package^6^ (v2.5.45-v.2.5.62). We further implemented a “joint” model fitter in HyPhy, which estimates a single set of MSS parameters from G>1 genes by maximum likelihood. The joint model optimizes α parameters (shared by all genes) for the MSS model by defining and maximizing the likelihood function on multiple genes, while holding gene-specific estimates of “nuisance” parameters, such as equilibrium codon frequencies, relative branch lengths (each gene receives a single multiplicative branch length scaler for the joint phase), nucleotide substitution biases, and ω at values obtained by fitting the standard MG94 model to each gene separately. Using gene-level “plug-in” estimates dramatically lowers estimation complexity and run times.

### Forward Population Genetic Simulations

To test that the method accurately identifies codon pairs under selection, a forward Wright-Fisher population genetic simulator was written (Pivirotto 2024). To begin, for each simulation an ancestral gene sequence of 300 codons was generated with codon frequencies based on a gene alignment randomly selected from the 1613 alignments used by Rahman *et al*.(Rahman et al. 2021). The ancestral gene sequence was copied to a population size of N=500 diploid individuals to found an ancestral population. The population was allowed to evolve to mutation-selection-drift equilibrium with two burn-in periods, with each generation including mutation and selection (without dominance) following a standard Wright-Fisher model (Ewens 2004). The first burn-in period proceeded for 200xN generations, a value that was determined empirically to be more than sufficient to reach a point when population mean fitness was stable for a wide range of mutation and selection parameters. The second burn-in period proceeded until all gene copies had a single common ancestor in the generation when the first burn-in ended. Following the burn-in period, the simulation continued for a preset number of generations with 10 population splitting events at set time points (Figure S1). Finally at the end of the simulation, an alignment was created by sampling a single, random gene copy from each of the 11 populations. Nonsynonymous mutations were all given a deleterious selection coefficient of 2Ns = -10 (*i.e.,s* = − 0. 01). The total depth and length of the tree was 10,000 and 64,000 generations respectively. The mutation rate per base position was 1. 6 × 10^−5^ per generation, which yielded approximately 1,850,000 neutral synonymous mutations and 230 neutral synonymous substitutions over the entire tree, for about 1.7 substitutions per effective synonymous site.

Synonymous selection was modeled by partitioning the codons for each amino acid into one or two neutral classes. In cases of one neutral class, codons within the neutral class were all neutral with respect to each other. In cases of two neutral classes, the codons in one class, selected arbitrarily, were favored over those in the other class, and changes within each class were treated as neutral. Mutations in the favored direction had a selection coefficient of s, while those in the reverse direction had a selection coefficient of -s. Sets of 100 simulated alignments were generated for values of 2Ns: 0, 0.5, 1.0, 1.5, 2.0, and 5.0. The assignment, selected or neutral, for each synonymous codon pair in each set of 100 genes, was inferred using a genetic algorithm search procedure implemented in HyPhy (see below). The data-generating (true) model included 25 selected codon pairs and 42 neutral pairs (Table S1).

### A genetic algorithm for learning synonymous substitutions from sequence alignments

Given a collection of **S**≥1 training alignments, and the desired number of synonymous rate classes (1 < **M** < N) we can infer (learn) how to partition all synonymous substitutions into M groups to maximize model fit. Because such an optimization involves exploring a large discrete space of models without natural ordering, we applied the CHC heuristic aggressive genetic algorithm approach (Eshelman 1991) which has shown to work well for evolutionary model selection (Kosakovsky Pond et al. 2007; Delport, Scheffler, Botha, et al. 2010), and discrete optimization (Kosakovsky Pond, Posada, Gravenor, Woelk, and Simon D. W. Frost 2006) previously. An evolutionary model is uniquely encoded by a D-dimensional vector (D = number of unique synonymous rates), where entry *i* is an integer in [0,M), indicating that rate *i* belongs to the corresponding rate class. To avoid model specification redundancy we enforce class-sortedness, *i.e.*, among all vectors that encode the same partition of rate classes we choose the one, where the first occurrence of class 0 precedes the first occurrence of class 1, etc. For example, (0,1,1,0,1) and (1,0,0,1,0) encode the same partition, but the former is selected as the “sorted” representative. The logic of the MSS-CHC algorithm is as follows, and key model parameters are enumerated in Table 3.

Step 1. Fit the standard MG94-REV model to each of the **S** input alignments to obtain branch lengths and the BIC score for the baseline or null model (all synonymous rates are equal). For computational efficiency, we fix branch lengths and nucleotide rates θ at their maximum likelihood estimates (MLEs) for the rest of the algorithm.

Step 2. Initialize **P** individuals (models) randomly, *i.e.*, generate **P** unique D-dimensional vectors encoding MSS models. For each model, *m*, evaluate its fitness *f*(*m*). This fitness is defined as the Bayesian Information Criterion (BIC) score for the fit of the model to all S training alignments. The sample size, needed for BIC computation, is defined as the total number of characters (sequences x sites) in all alignments. Record the best current fitness *F* score (lowest BIC), and the corresponding most fit individual *Fm*

Step 3. Create the next generation through recombination and elitist selection. To do so, generate **P** offspring individuals, by randomly selecting two parents from the current generation, and performing free recombination on them: each entry of the child vector has a 50-50 probability of coming from either parent. The parent and offspring populations are combined and top **P** most fit individuals are retained.

Step 4. Decide if the population is too inbred and it needs to be mutagenized. This will occur if at least one of the following two conditions is met. First, the relative fitness range (relative difference between the most and the least fit model) is too low (*f_rel_*). Second, the number of consecutive generations during which Step 3 failed to introduce any new models is too high *f_stagnant_*. To mutagenize the population, each individual, except for the most fit one, is randomly mutated with probability μ of changing each element of the model vector.

Step 5. Check the termination criterion: does the number of consecutive generations without a sufficient (δ_*BIC*_) BIC improvement exceed *G_conv_*? If the termination criterion is met exit, otherwise go to Step 3.

This search procedure is implemented in the HyPhy script MSS-GA.bf. The output of an MSS-CHC model search is a JSON file which records every model considered by the algorithm (*i.e.,* its D-dimensional encoding), the corresponding fitness (BIC score), and MLEs for synonymous rate parameters (α). The final model is derived from the MSS-CHC run using model averaging (Kosakovsky Pond, Posada, Gravenor, Woelk, and Simon D.W. Frost 2006). Specifically, each model is given an Akaike Weight between 0 and 1, by normalizing its BIC score as follows: *w*(*m*) = *exp*((*min_models_BIC*(*m*) − *BIC*(*m*))/2)/*K* with K chosen so that add up to 1. Then, the rate estimates from each model are ordered from smallest (“selected”) to largest (“neutral”), with intermediate values labeled as “INT 1”, “INT 2” etc, based on their rank, and a categorical vector is derived. For example, if the model vector was (0,1,1,2,0,2) with α_0_ = 1, α_1_ = 0. 8, α = 1. 4, then the resulting categorical model vector would be `(INT 1, SELECTED, SELECTED, NEUTRAL, INT 1, NEUTRAL)`. Finally, for each of the **D** elements (rates) of the final model we compute cumulative Akaike weights that accrued to each category for that rate. A rate assignment is considered to be unambiguous, if a particular category (SELECTED, NEUTRAL, etc) garners 0.90 or more of the total weight, otherwise, it is considered ambiguous.

### Empirical data collection and preparation

We used gene alignments from twelve *Drosophila* species, including *D. melanogaster*, for which there is existing strong evidence of natural selection on synonymous sites (Shields et al. 1988; Akashi 1994; Hey and Kliman 2002). The other species included in our analysis (*D. erecta, D. rhopaloa, D. elegans, D. santomea, D. simulans, D. mauritiana, D. subpulchrella, D. sechellia, D. suzukii, D. takahashii,* and *D. yakuba)* also have available genome assemblies and similar patterns of codon usage as *D. melanogaster* (Supplemental File D12_codon_usage.zip). Soft-masked genomes and the *D. melanogaster* reference genome were downloaded from the genomes database of the National Center for Biotechnology Information (NCBI) (Supplemental File Drosophila_genomes_aligned.txt). A whole genome alignment with respect to the *D melanogaster* genome was generated using cactus (Armstrong et al. 2020) and the hal2mafMP.py script, and the phylogenetic tree of Suvorov *et al* (Suvorov et al. 2022).

To generate gene alignments we mapped the location of *D. melanogaster* gene coding sequences (CDSs) in the reference genome sequence, and then built gene alignments from the whole genome alignment. A species’ gene sequence was rejected from the alignment if any sequence fragments were not in the same order as the reference, or if any exons were not in the same order as the reference or not on the same strand, or if the codon frame of an exon did not match that of the reference, or if there were any stop codons. Gene alignments were rejected if their nucleotide length was not a multiple of 3 and if there were fewer than 4 species represented. These criteria yielded alignments for 11,844 genes.

#### Drosophila polymorphism analysis

If the inferred substitution rate of a codon pair reflects the extent of (purifying) selection occurring, then codon pairs with higher relative substitution rates should be closer to selective neutrality. A straightforward prediction of this hypothesis is that polymorphism densities taken from population genomic data for each of the 67 synonymous codon pairs should be positively correlated with substitution rates. In other words, codon pairs that have higher substitution rates should also have higher counts of segregating SNPs in a natural population. To test this prediction we counted synonymous SNPs in a *D. melanogaster* population genomic sample of 197 genomes from Zambia (Lack et al. 2015) for each of the 67 synonymous codon pairs. Then for each gene we estimated the density of each codon pair, by dividing the SNP count for a pair by the count of the corresponding amino acids.

#### Quantifying tRNA abundance

We obtained tRNA abundance data for *Drosophila melanogaster* as raw transcript counts per tRNA gene from Behrens *et al*.(Behrens et al. 2021) (GEO accession number GSE152621). We normalized the reads to counts per million (CPM), but did not also normalize by gene length because tRNA genes are all roughly the same length (76 - 90 bp)(Sharp et al. 1985). From these data, we had the tRNA abundances for 45 out of 61 codons. Some tRNA modifications, specifically Inosine and Queuosine, expand the decoding ability of tRNA isoacceptors.

When the anticodon wobble base A is biochemically modified to inosine (a Guanine analog)(Torres et al. 2015), isoacceptor recognition expands to include codons ending in U, C, or A. In eukaryotes, Inosine modifications are found on the wobble bases of the anticodons AGC (Ala), AGG (Pro), ACG (Arg), AGA (Ser), AAG (Leu), AAC (Val), AAT (Ile), and AGT (Thr). We added the abundance of each modified tRNA to the pool of tRNAs that could decode synonymous codons ending in A or C. Because the anticodon wobble base is A, its abundance already counts towards the cognate U-ending codon pools.

When the anticodon wobble base G for tRNAs that decode His, Asp, Asn, and Tyr (Chiari et al. 2010) is modified to a Queuosine, the isoacceptors’ decoding capacity expands to include synonymous U-ending codons in addition to the cognate C-ending codons. We therefore added the abundance of each modified tRNA to the pool of synonymous tRNAs that could decode cognate U-ending codons.

After accounting for the expanded decoding capabilities of Queuosine- and Inosine-modified tRNAs, there were six codons where with multiple possible tRNA decoders: UUU (Phe), UGU (Cys), CGG (Arg), AGU (Ser), GGU (Gly), and GGG (Gly). For these codons, we assigned them the abundance of the tRNA most likely to decode them based on Crick’s wobble rules (Crick 1966). Specifically, we assumed that twofold-redundant amino acids decode both of their codons using a single tRNA isoacceptor that recognizes U- and C-ending codons. We gave UUU (Phe) an equal tRNA abundance to UUC (Phe), gave AGU (Ser) an equal tRNA abundance to AGC (Ser), and gave UGU (Cys) an equal tRNA abundance to UGC (Cys). We deduced that the remaining amino acids (Arg and Gly) were not decoded by one tRNA that recognizes U, C, or A in the wobble position and one tRNA that only recognizes G, as neither amino acid had any tRNA abundance data for G-ending codons. We also deduced that Arg codons beginning with C could not be served by the same tRNA as Arg codons beginning with A, since the first codon position does not wobble. Therefore, we assumed that CGG (Arg) must be decoded by a single tRNA isoacceptor that recognizes G- and A-ending codons, and gave CGG (Arg) and equal tRNA abundance to CGA (Arg). We also assumed that Glycine had one tRNA isoacceptor that recognizes G- and A-ending codons (giving GGG (Gly) the same abundance as GGA (Gly)), and another isoacceptor that recognizes U- and C-ending codons (giving GGU (Gly) the same abundance as GGC (Gly)).

We made the necessary simplifying assumptions that (i) all tRNAs capable of being modified are in fact modified, (ii) modification is present in all tissues across all life stages, and (iii) modified tRNAs are equally capable of binding to all codons they can recognize. The true biology is undoubtedly more complex (for example, the level of queuosine modification is dependent on the availability of dietary queuine (Zhang et al. 2022)), but teasing apart the exact tRNA pools across different conditions in a living animal is an open research question beyond the scope of this manuscript.

## Results

### Testing MSS by forward simulation

Like the MG94 model (Muse and Gaut 1994) it is based on, MSS models (including SynREVCodon) implement a model of codon substitutions as a reversible time-homogeneous Markov process with independent sites. MSS models method estimate rate parameters for synonymous codon pairs, with the implicit assumption that for a given codon pair the substitution rate in each direction is the same. However if one codon is favored over the other then the mean waiting time to the favored codon will be shorter on average than the mean waiting time to the disfavored codon, violating a key MSS modeling assumption. We assessed MSS model performance under a more biologically realistic “mis-specificed” model using a forward population genetic simulation without the reversibility assumption, and with directly parameterized selective coefficients on a predefined set of codon pairs.

Forward population genetic simulations model the evolution of a population(s) over time by simulating the reproduction of individuals or genes as a function of their fitness. To generate data suitable for testing MSS models, a run of our simulator begins with a population that evolves to the point of mutation-drift-selection equilibrium, and then continues to evolve with periodic population splitting events to generate a final series of 11 populations from which individual sequences were then sampled.

To simulate data with a single selection coefficient on a subset of synonymous codons, we specified pairs of one-step reachable codons that were neutral with respect to each other within each redundantly encoded amino acid, then assigned a selection coefficient to apply to pairs of codons that fell within separate neutral pair sets. We executed 100 simulations for varying levels of selection, parameterized through *2Ns* (Table 4), each under a model with 25 selected codon pairs and 42 neutral pairs; these pairs were selected *a priori*. We next ran the genetic algorithm search procedure to infer neural and selected codon pair sets from sequence data. This classifier partitions all 67 codon pairs into non-overlapping sets, and assigns a numerical confidence of set membership (between 0 and 1) to each pair.

**Table 1:** Instantaneous substitution rates from one sense codon (x) to another (y)

| One nucleotide? | Synonymous? | Substitution rate | Example |
| --- | --- | --- | --- |
| Yes | Yes | $\alpha(x, y) \theta(x, y) \pi(x, y)$ | $AG\bar{C} \rightarrow AG\bar{T} : \alpha_{AGC:AGT} \theta_{CT} \pi_T^3$ |
| Yes | No | $\beta(x, y) \theta(x, y) \pi(x, y)$ | $A\bar{G}\bar{C} \rightarrow A\bar{C}\bar{C} : \beta \theta_{CG} \pi_C^2$ |
| No | * | 0 | $\bar{T}\bar{C}\bar{C} \rightarrow \bar{A}\bar{G}\bar{C} : 0$ |

**Table 2:** The way that different MSS models model variation among synonymous substitution rates, α (*x*, *y*).

| Model | Functional form | Parameters in addition to basic MG94 |
| --- | --- | --- |
| Standard codon | $\alpha(x, y) := 1$ | **0** |
| *A priori* | $\alpha(x, y) = \alpha_i, i = 1..K$ | **K - 1, $1 < K < N$**, where **N** is the number of amino-acids that have synonymous codons that can be exchanged by a single nucleotide substitution, e.g, **K=2** in Rahman *et al* |
| Genetic algorithm<br>(GA) | $\alpha(x, y) = \alpha_{<i>, i = 1..K$ | **K - 1, $1 < K < N$**. Same as above, except the allocation of specific (x,y) pairs to the i-th rate class is not fixed *a priori* but inferred using a heuristic discrete space search. |
| SynREV | $\alpha(x, y) = \alpha_A$<br>A is the aminoacid encoded by x and y | **N - 1 (18 for the Universal genetic code)** |
| SynREVCodon | $\alpha(x, y) = \alpha_{xy} = \alpha_{yx}$ | **P - 1**, where **P** is the number of synonymous codon pairs within one nucleotide substitution of one another (**67** for the Universal genetic code) |

**Table 3:**
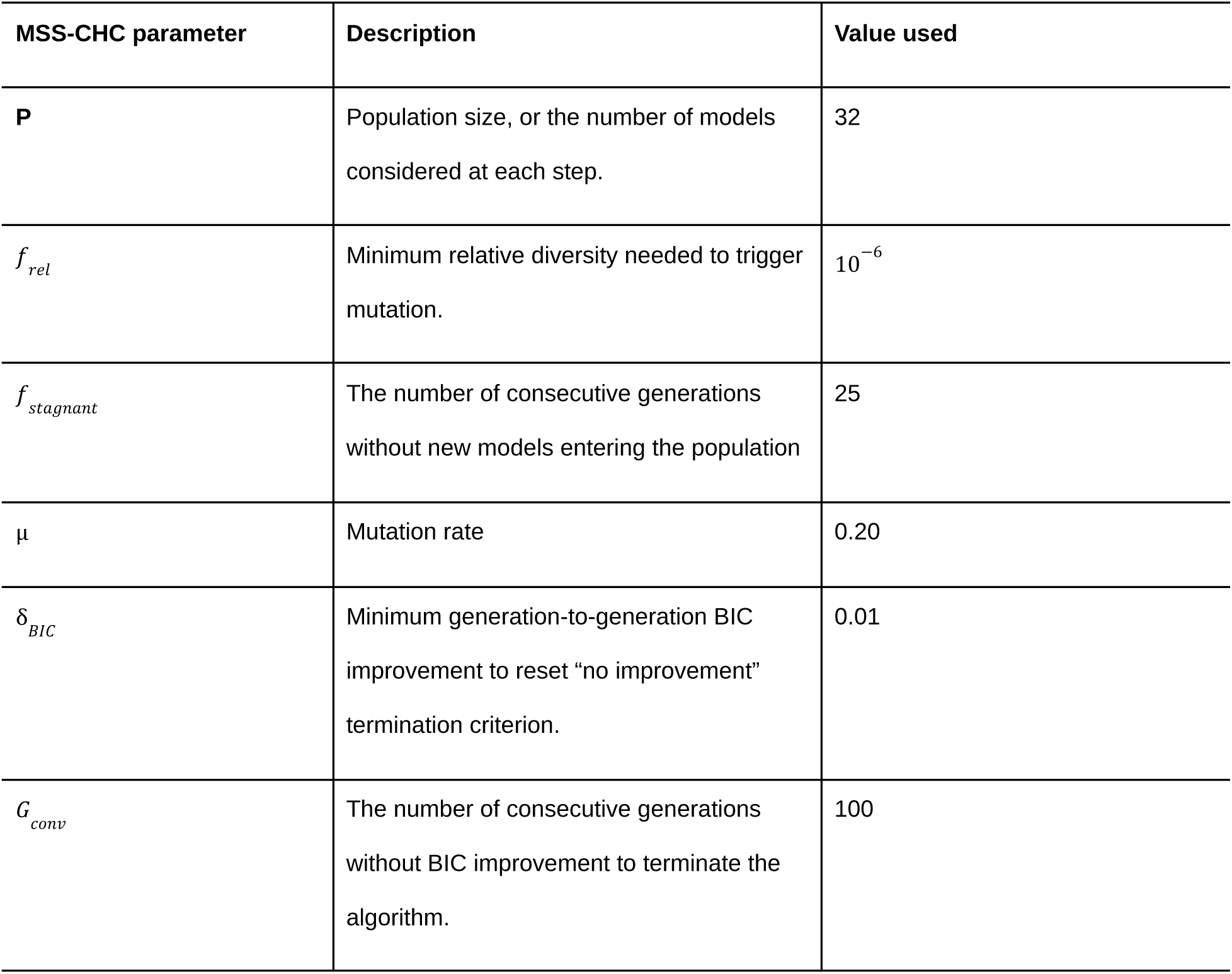
Key parameters of the CHC genetic algorithm for model selection.

**Table 4:** The results of genetic algorithm model selection applied to N=100 alignments obtained using forward simulations for different strengths of selection (0: neutral, -5 : strong purifying selection). ΔBIC reports the improvement by the best model found by the genetic algorithm search compared to the baseline MG94 model. The 2×2 contingency table of simulated/inferred classifications for each codon pair, the corresponding p-value for association, and the Matthews Correlation Coefficient (MCC) are provided. For classification purposes, we place each codon pair in the Selected/Neutral class based on which received more model averaged support. To quantify the degree of assignment uncertainty, we also tabulate how many of the 67 rates were “ambiguous”, i.e., had less than 0.9 support for the class assignment.

| **Selection**<br>**coefficient**<br>**2Ns** | **ΔBIC** | **MCC** | **Fisher**<br>**Exact test**<br>**p-value** | **Inferred**<br>**codon**<br>**classes** | **Simulated (true)**<br>**codon classes** |  | **Ambiguously**<br>**assigned pairs** |
| --- | --- | --- | --- | --- | --- | --- | --- |
|  |  |  |  |  | **Selected**<br>**(25)** | **Neutral**<br>**(42)** |  |
| 0 | 152.8 | -0.25 | 0.06 | Selected (45) | **12** | 32 | 10 |
|  |  |  |  | Neutral (22) | 12 | **10** |  |
| -0.5 | 192.6 | 0.093 | 0.54 | Selected (53) | **21** | 32 | 7 |
|  |  |  |  | Neutral (14) | 4 | **10** |  |
| -1 | 282.5 | 0.51 | 4.02e-05 | Selected (29) | **19** | 10 | 6 |
|  |  |  |  | Neutral (38) | 6 | **32** |  |
| -1.5 | 675.0 | 0.65 | 1.7e-7 | Selected (26) | **20** | 6 | 4 |
|  |  |  |  | Neutral (41) | 5 | **36** |  |
| -2 | 1630.4 | 0.81 | 2.07e-11 | Selected (23) | **21** | 2 | 3 |
|  |  |  |  | Neutral (44) | 4 | **40** |  |
| -5 | 17974.7 | 0.97 | 2.6e-17 | Selected (24) | **24** | 0 | 0 |
|  |  |  |  | Neutral(43) | 1 | **42** |  |

The model selection procedure does progressively better as the strength of negative selection is increased. This is evidenced by the progressively improving model fit (ΔBIC), better agreement with the ground truth (MCC, Fisher’s Exact test p-value), and the contingency table itself. For strong selective regimes, the inference procedure is highly accurate, and starting at a relatively weak selective level (2Ns = -1), there’s significant predictive value in the inferred codon classes. Thus, despite being provided with data simulated under a cardinally different model, MSS Markov models are able to infer the set of neutral codon pairs with high accuracy, assuming that selection acting on the non-neutral codon pairs is sufficiently strong. The genetic algorithm procedure remains sensitive to model misspecification, since it still finds statistically significant improvement for an MSS model compared to the no selection (Ns = 0) model, based on ΔBIC. Because forward-simulated data is generated under a model that is different from the time-reversible Markov model assumed by the MSS, apparent improvement in fit may be due to phenomenological loading (Jones et al. 2018). As a separate exercise we also ran GA model selection on parametrically simulated under the properly specified model (MG94). When null data (100 alignments per run, 25 replicates) are simulated parametrically under the expected MG94 model, the genetic algorithm procedure correctly selected the null (no synonymous rate variation) model in 22/25 simulated cases, and in the remaining 3 cases had ΔBIC ≤ 2 (detailed data not shown, see Supplementary Information for references).

### Synonymous substitution rates in *Drosophila* genes

#### Description and information content of the data

We analyzed 11,844 gene alignments from *Drosophila* containing a median of 12 (range 4-23) sequences with a median of 411 (range 21 - 8933) codons, and median tree length of 0.48 (range 0.017 to 2.0) substitutions per nucleotide site (Figure S1). To gain some intuition of data content in these genes that is relevant to MSS inference, we mapped codon substitution to branches using joint maximum likelihood ancestral reconstruction methods implemented in SLAC (Kosakovsky Pond and Frost 2005) and counted approximately how many single-nucleotide synonymous substitutions (SNSS) occurred in *Drosophila* genes. The entire collection of genes was estimated to include ∼4.12M SNSS, and this can be viewed as an “order-of-magnitude” approximation of the sample size available for MSS inference. Per gene, there was a median of 254 (range 0-5762) SNSS, and as expected, this number was positively correlated with gene length and divergence (Figure S2). Among 67 types of SNSS available in the universal genetic code (*e.g*. AAA↔AAG, a substitution between Lysine codons), there was an order of magnitude difference between how frequent individual substitutions were (Figure 1, Table S2), implying that some rates might be easier to infer than others, and some rate estimates might be less noisy. Furthermore, for a fixed amino-acid with more than one SNSS, there was generally a significant difference between the least common and the most common pair of synonymous codons being substituted. For example, there was a ∼6-fold difference in raw counts between the most (CTG↔TTG, ∼160K) and least (CTA↔TTA, ∼25K) common SNSS among Leucine codons, even though the same pair of nucleotides are involved (C,T). This disparity is relevant in the context of codon-level MSS models, *i.e*., when individual codon pairs coding for the same amino-acid may have unequal relative substitution rates. Relatively rare substitutions, *i.e.,* those present at small count numbers (*e.g.,* TCA↔TCT for Serine, having a mean of 1.21 per alignment), or those found only in a minority of datasets (*e.g*., TCA↔TCT is found in 5808/11844 alignments) can have larger sampling noise, and may be inferred less accurately. SNSS count distributions have heavy tails (Figure 1), with a significant number of genes where substitution counts for a pair significantly exceed the mean. This pattern will help explain why rate estimates under the MSS models may also be expected to have heavy tails, with large estimates for individual genes.

**Figure 1.**
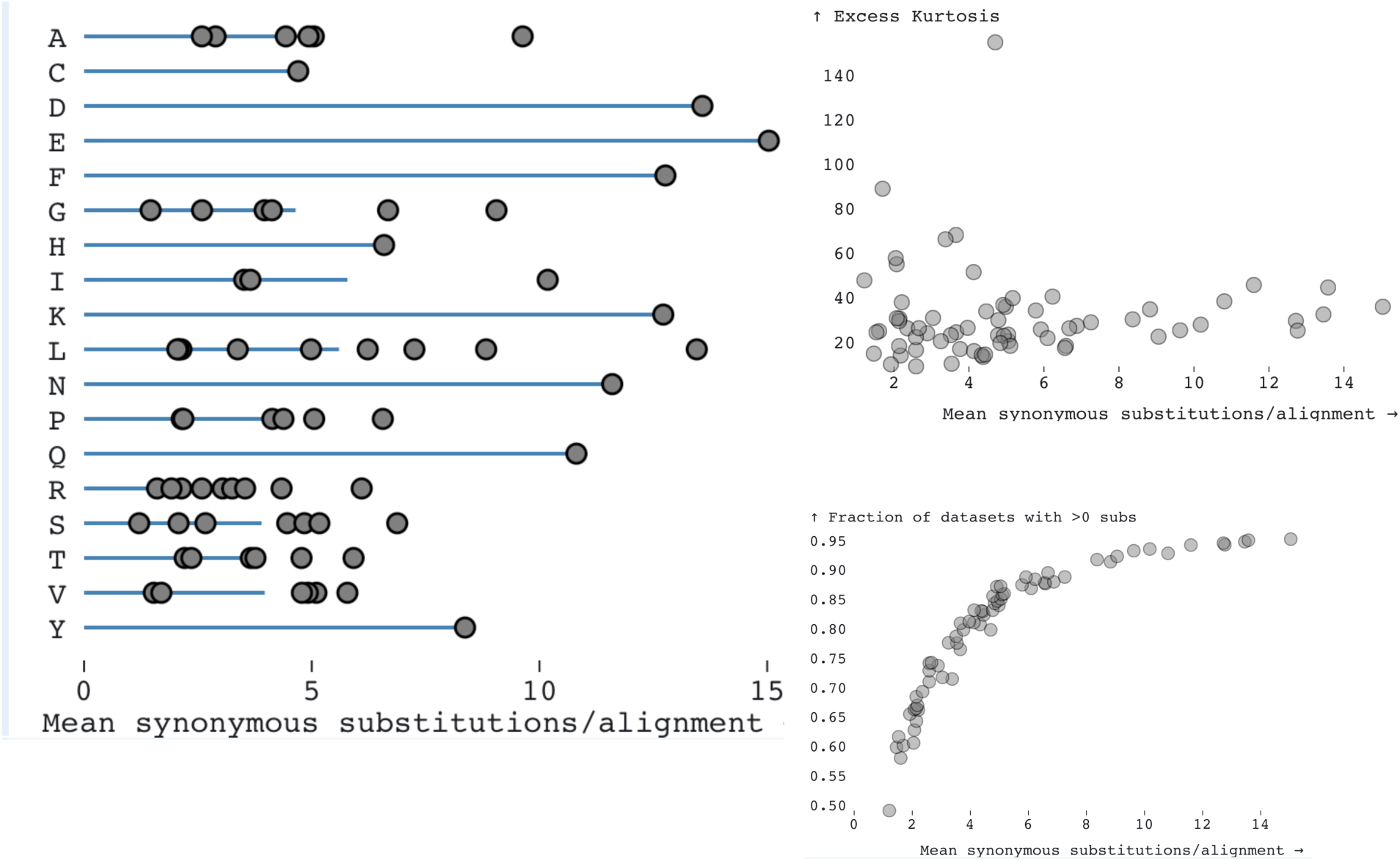
*Left*: the average (per alignment) number of SNSS for each of the 67 codon pairs in the Universal genetic code (grey dots) inferred from the *Drosophila* gene alignments. Blue lines demarcate averages over individual codon pairs for each amino acid. *Right (top):* the distribution of the mean number of SNSS codon-pairs per alignments vs the excess kurtosis shows that all of the MSS rates have a large number of observations in the tail end of the rate distributions. *Right (bottom):* the distribution of mean SNSS codon-pairs per alignment vs the number of alignments where >0 of substitutions for the specific codon-pair are present.

#### Estimating synonymous substitution rates using maximum likelihood

The most general model in the MSS class is SynREVCodon which directly estimates the relative substitution rate for each of the 67 synonymous codon pairs, α_*i*_. SynREVCodon has 66 identifiable relative synonymous rate (RSR) parameters. Because the mean of the synonymous substitution rates is confounded with branch lengths (only products of α and t can be estimated), we impose the constraint Σ α_*i*_ / 67 = 1. We also consider a simpler model, SynREV, which is nested in SynREVCodon and sets all the pairwise rates for codons encoding the same amino-acid to be the same. This results in 18 amino-acid level codon parameters (methionine and tryptophan each have a single corresponding codon).

We fit MSS models to the collection of *Drosophila* alignments in two different ways. The first is the fully independent model (FIM), where each gene has its own set of synonymous substitution rates α_*i*_, along with other parameters (branch lengths, nucleotide rate biases, codon frequencies, and the non-synonymous rates). The second is the fully shared model (FSM), where the same set of α parameters is inferred jointly from all genes, while all other parameters are estimated independently for each gene. More precisely, we estimate branch lengths, nucleotide rate biases, codon frequencies, and the non-synonymous rates using the standard MG94xREV model for each gene, and then hold those at their point estimates while fitting the MSS rates jointly, and also estimating an overall tree length scaling parameter for each gene (proportional branch lengths are maintained from gene level fits). These approximations make the multi-gene fit computationally feasible.

The FIM model is expected to produce noisy gene-level estimates because parameter-rich models are fitted to relatively small alignments, but it is useful for quantifying the extent of gene-level variability in rates. The FSM model is expected, on the other hand, to yield “average” estimates with low variance, but has no capacity to account for gene-to-gene heterogeneity. Because the standard MG94 model (α ≡ 1) is nested within the MSS (SynREV and SynREVCodon) model, a standard likelihood ratio test (LRT) with 17 (SynREV) or 67 (SynREVCodon) degrees of freedom can be used to test the hypothesis that some synonymous substitution rates are different, gene-by-gene, and genome-wide.

We fitted the nested hierarchy of models MG94xREV ⊂ SynREV ⊂ SynREVCodon to each of the 11,844 gene alignments, and compared them to each other via likelihood ratio tests (Table 5). Despite the relatively small sizes of individual alignments, the standard codon model can be rejected in favor of the parameter-rich MSS models for a substantial fraction (SynREV) or an outright majority (SynREVCodon) of the alignments, *e.g.*, based on false discovery rates. The family-wise error rate correction (Holm-Bonferroni) is much more conservative. Among the two MSS models, SynREV can be rejected in favor of SynREVCodon on the majority of the tested alignments.

**Table 5.**
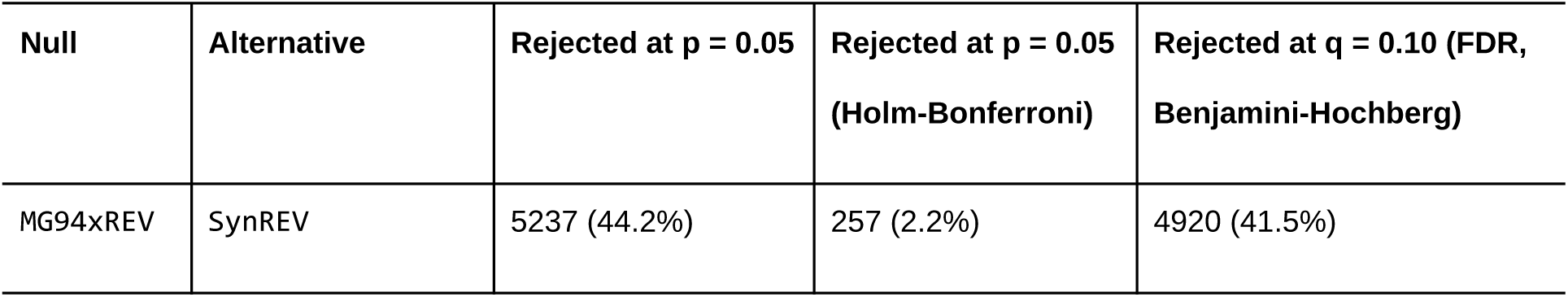
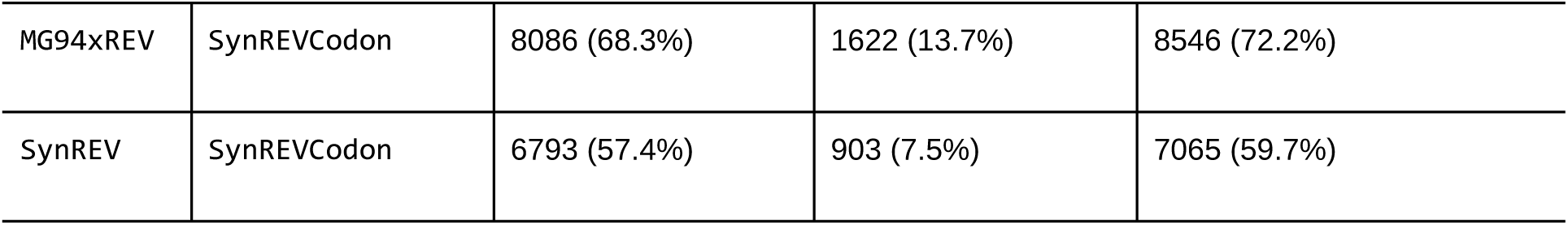
Model comparison results using likelihood ratio tests on individual *Drosophila* gene alignments.

**Table 6.** Fit improvements when comparing MG94xREV to SynREV and SynREVCodon models fitted jointly to subsets of multiple alignments. Means and ranges refer to the five random subsets of genes for a given batch size.

| # | LRT p-value vs MG94 p-value | ΔAIC / alignment |
| --- | --- | --- |

| genes | Mean (range) |  | Mean (range) |  |
| --- | --- | --- | --- | --- |
|  | SynREV | SynREVCodon | SynREV | SynREVCodon |
| **4** | 0.121(0.000, 0.599) | 0.023(0.000, 0.114) | 0.369(-4.762, 2.415) | -3.973(-13.203, -0.518) |
| **8** | 0.012(0.000, 0.038) | 0.000(0.000, 0.002) | 0.561(-0.672, 1.683) | 0.138(-3.722, 4.881) |
| **16** | 0.000(0.000, 0.000) | 0.000(0.000, 0.000) | 2.550(1.261, 4.521) | 5.198(2.331, 10.588) |
| **32** | 0.000(0.000, 0.000) | 0.000(0.000, 0.000) | 3.169(1.576, 4.170) | 8.249(5.252, 12.248) |
| **64** | 0.000(0.000, 0.000) | 0.000(0.000, 0.000) | 3.469(2.615, 3.882) | 9.435(7.614, 10.026) |
| **128** | 0.000(0.000, 0.000) | 0.000(0.000, 0.000) | 3.819(3.427, 4.333) | 10.060(9.031, 10.680) |
| **256** | 0.000(0.000, 0.000) | 0.000(0.000, 0.000) | 3.843(3.628, 4.181) | 10.092(9.369, 11.070) |
| **512** | 0.000(0.000, 0.000) | 0.000(0.000, 0.000) | 3.608(3.338, 3.852) | 10.100(9.285, 10.624) |
| **1024** | 0.000(0.000, 0.000) | 0.000(0.000, 0.000) | 3.799(3.678, 3.907) | 10.413(10.239, 10.735) |
| **2048** | 0.000(0.000, 0.000) | 0.000(0.000, 0.000) | 3.718(3.617, 3.771) | 10.261(10.075, 10.380) |

Ranking the rates inferred by SynREVCodon by their median (across genes) estimates reveals a clear gradient (Figure 2), with TTC:TTT (Phenylalanine) as the fastest pair, and GGC:GGG (Glycine) as the slowest pair. Because MSS models are quite parameter rich, and individual alignments are not large, many rate estimates are exactly 0 (*e.g.*, no corresponding substitutions have occurred), and many other relative rate estimates are large, resulting in bimodal distributions. Similarly, there is a gradient of median relative amino-acid rates inferred by the SynREV model (Figure 3), with Alanine being the slowest, and Phenylalanine – the fastest. In the following sections we will show that some of this variation is attributable to cognate tRNA abundance, and is significantly correlated with independent measures of selection, namely population level polymorphism data.

**Figure 2.**
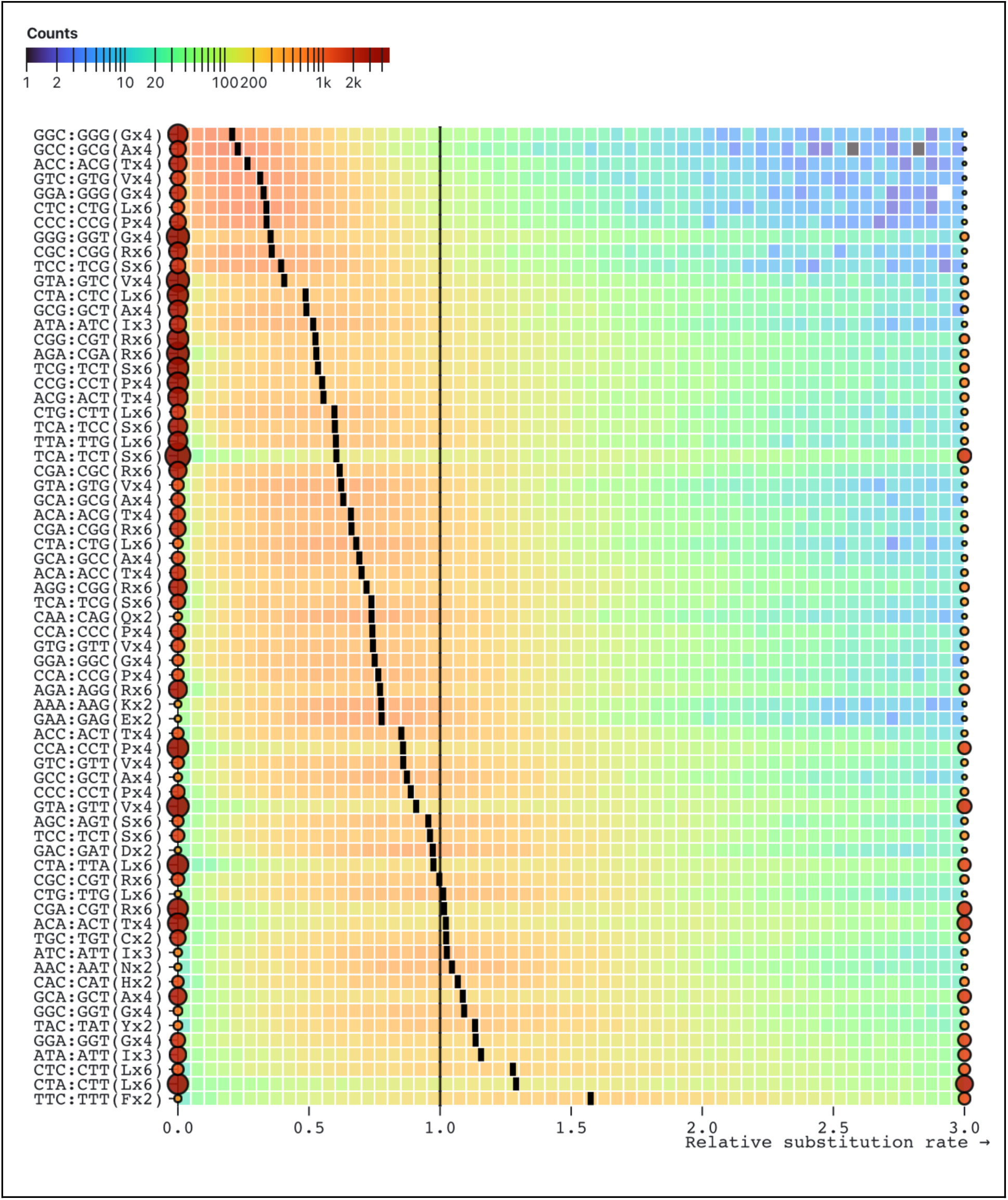
Maximum likelihood point estimates for relative rates of each of the 67 pairs of synonymous substitutions in the SynREVCodon model. The rates are sorted by the median of the corresponding distribution (black tick). Rates that are exactly zero or those that are ≥3 are shown as circles, whose size and color reflect the number of alignments contributing to the corresponding estimates. The heatmap for each pair is generated using only the rates that are in (0,3).

**Figure 3.**
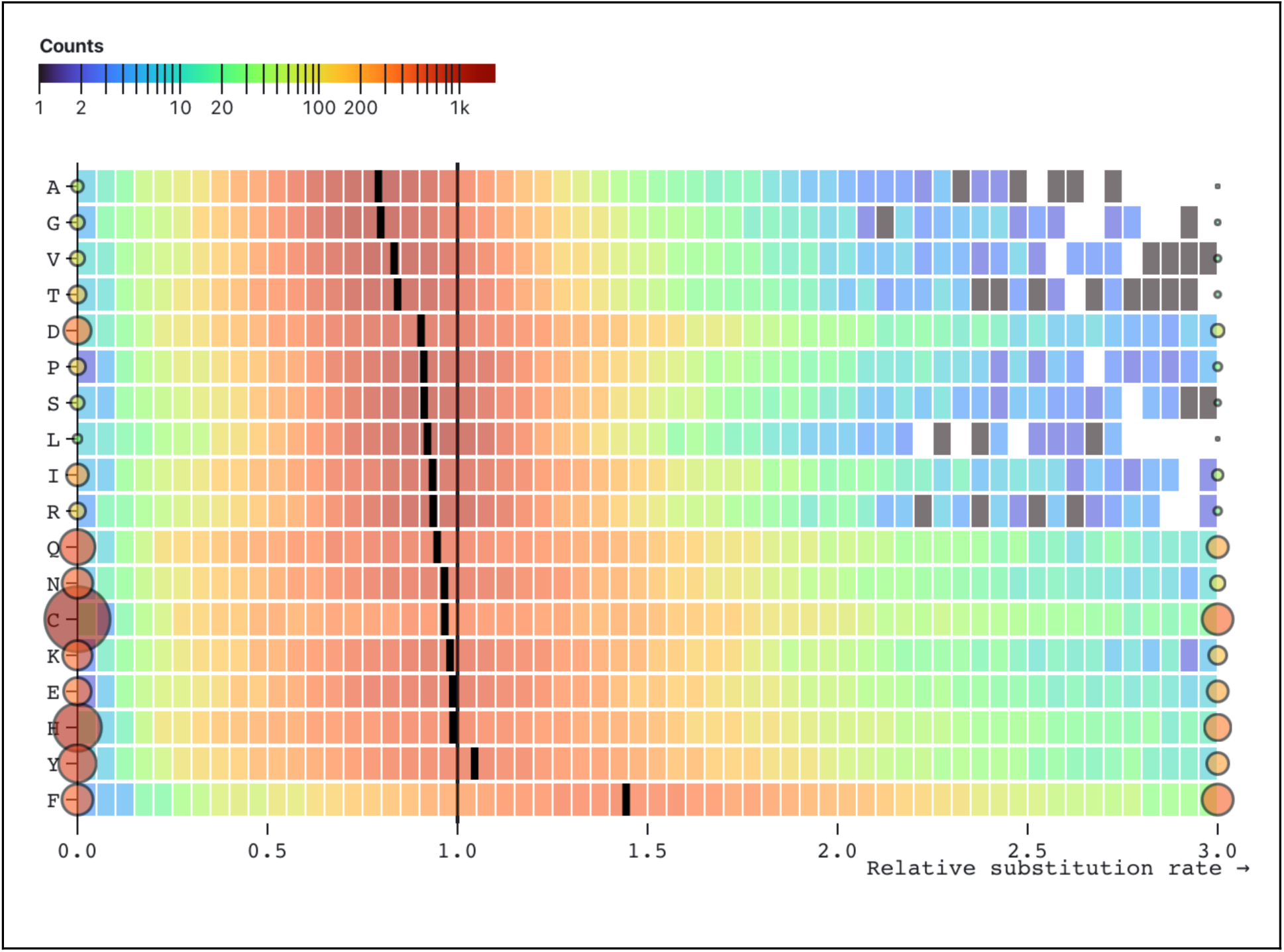
Maximum likelihood point estimates for relative rates of each of the 18 amino-acid level synonymous substitutions in the SynREV model. The rates are sorted by the median of the corresponding distribution (black tick). Rates that are exactly zero or those that are ≥3 are shown as circles, whose size and color reflect the number of alignments contributing to the corresponding estimates. The heatmap for each pair is generated using only the rates that are in (0,3).

For the nine amino-acids that have more than a single synonymous codon pair exchangeable by a single nucleotide substitution (Figure 4), the rates for individual codon pairs are often quite different and not well represented by a single (amino-acid level) rate. For example, for Leucine there are nine synonymous codon pairs exchangeable by a single nucleotide substitution. The medians of the corresponding rate estimates range from 0.34 (CTC:CTG conserved) to 1.29 (CTA:CTT neutral), while the median amino-acid level rate is 0.92. This observation, combined with the high rate of rejection of SynREV in favor of SynREVCodon (Table 5), provides compelling evidence that model variation in relative substitution rates should be modeled on the level of individual codon pairs.

**Figure 4.**
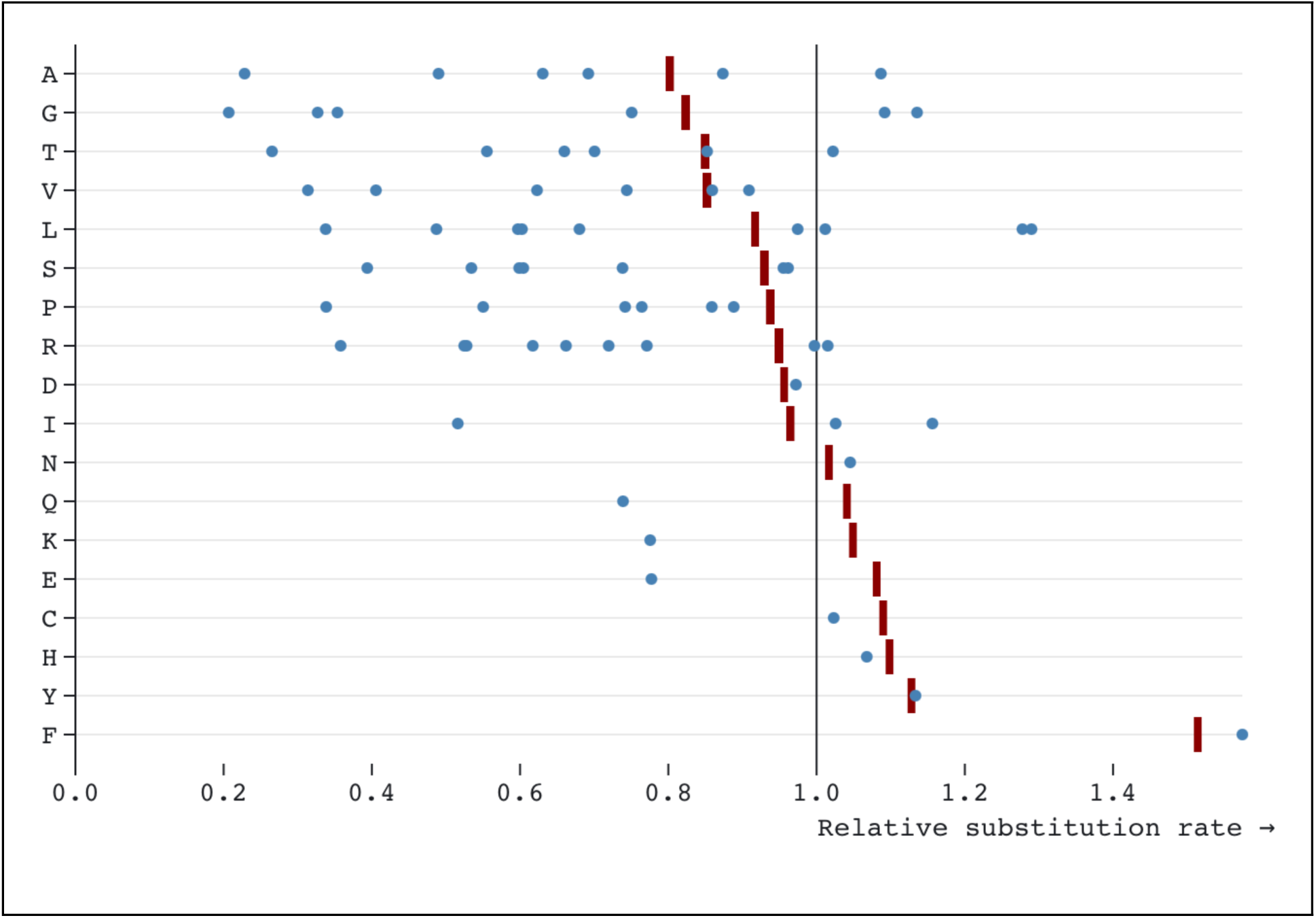
Median (over genes) estimates of relative substitution rates from SynREV (red ticks) and SynREVCodon (blue dots) models. For amino-acids with multiple pairs of synonymous substitutions, individual codon pairs are dispersed around the corresponding amino-acid level estimates.

### Codon pair substitution rates show strong correlation with polymorphism levels

For each pair of synonymous codons that differ by a single nucleotide, SynREVCodon provides an estimate of the corresponding relative rate of substitution. It does so while accounting for some mutational biases that give rise to uneven base composition, as well as the single-step rates from those two codons to other synonymous and nonsynonymous states. Natural selection operating on synonymous codon pairs is a plausible biological process contributing to inferred rate variation, although the complexity of the actual biological process and the unavoidable crudeness of substitution models make direct attribution unadvisable (Jones et al. 2018). Nonetheless, neutral codon pairs are expected to exhibit the highest relative substitution rates. For pairs where one codon is favored over another, the substitution process would alternate between relatively rapid substitution, from unfavored to favored, followed by a much slower rate of reversion. However, the average rate of selected synonymous codon pairs is expected to be lower than for neutral codon pairs, because the waiting time for the unfavored mutations is much longer than for the favored or neutral mutations (Kimura 1962; Rahman et al. 2021).

To ascertain how well the variation in codon pair rates reflects differences in selection, we turned to a corollary prediction from population genetics: alleles under directional selection will tend to be lost or fixed relatively rapidly relative to the neutral alleles. In other words, codon pairs that experience selection should have lower rates of polymorphism within a natural population, than should neutral codon pairs.

We estimated mean single nucleotide polymorphism (SNP) density across genes for each of the codon pairs, as observed in a collection of 197 genomes from Zambia, and compared them against the median of the corresponding substitution rate estimated from individual genes by SynREVCodon model (Figure 5). The linear correlation coefficient between the two quantities is 0.52 (corrected R^2^ of 0.26). Spearman rank-based correlation is similarly high (⍴=0.48) and significant (p = 0.002).

**Figure 5.**
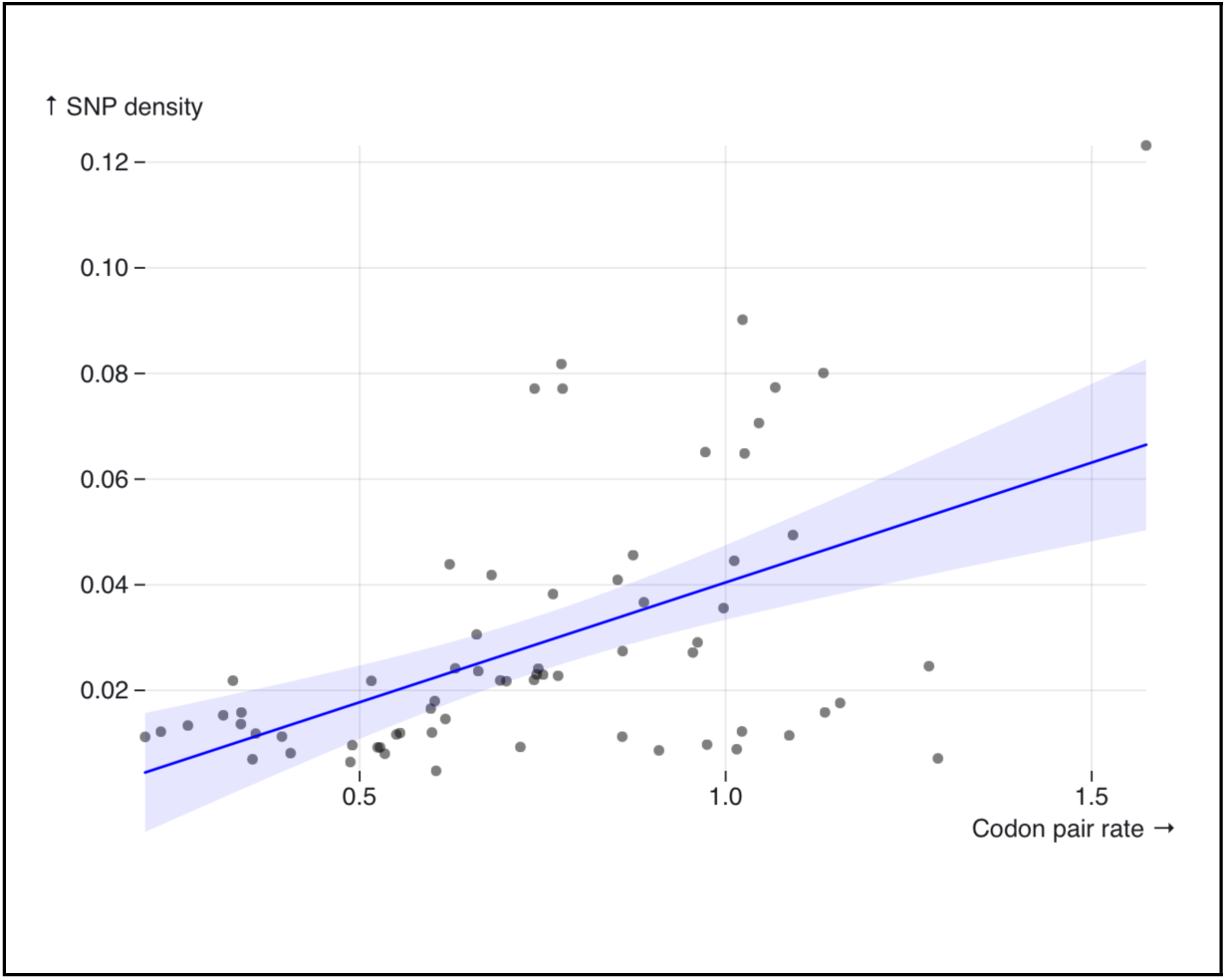
There is a positive correlation (Spearman’s ⍴ = 0.48) between the median inferred synonymous substitution rates, and the mean SNP density observed in a collection of 197 *Drosophila* genomes from Zambia. SNP density is an independent measure of selection pressure, as selection pressure to maintain a preferred codon decreases the SNP density for that codon. Correspondingly, the synonymous substitution rate for that codon will also be lower.

As the two data sets, SNP density in a natural population and species alignments used for SynREVCodon, are quite independent of each other, the finding of a strong correlation supports the claim that codon pairs with high estimated substitution rates are under less selection than those with lower estimated substitution rates.

### Codon pair substitution rates correlate with tRNA abundance

Researchers have long hypothesized that codon usage is under selection to conform with tRNA abundance, as a means of ensuring translational efficiency (Ikemura 1981; Kanaya et al. 2001). Compelling evidence for this hypothesis comes from the observation that codon usage bias is inversely correlated with gene expression: highly expressed genes almost exclusively use synonymous codons served by the most abundant tRNAs. We therefore assume that substitutions between a codon served by a highly abundant tRNA and a codon served by a less abundant tRNA would be disfavored, and that purifying selection would reduce the rate of such substitutions. To examine our assumption, we obtained and normalized tRNA transcript counts from Behrens *et al* (Behrens et al. 2021). We then examined tRNA modifications that increase the decoding capacity of individual tRNA isoacceptors. For each tRNA that was modified to recognize an additional codon, we added the modified tRNA’s abundance to the pool of tRNAs capable of decoding that codon. For codons where the decoder tRNA was ambiguous, we used Crick’s wobble pairing rules to predict which tRNA decoded which codon (Crick 1966).

We calculated the ratios in tRNA abundance between codons in a pair. Though SynREVCodon rates are symmetric (*e.g.*, the rate of the Phenylalanine substitution TTC > TTT is equivalent to the rate TTT > TTC), the abundance ratio is sensitive to the order in which abundances are compared (*e.g.,* if one tRNA pool is four times as abundant as the other, the abundance ratio may be 4/1 or ¼). To enforce consistency among the abundance ratios, we chose to always measure the abundance of the more common tRNA decoder pool compared to the less common one (*e.g.,* an abundance ratio of 4/1). Doing so allowed us to maintain a consistent interpretation of the relationship between substitution rates and abundance ratio magnitudes. We found a significant negative correlation between the median synonymous substitution rates inferred by SynREVCodon and the tRNA abundance ratios (rank-based Spearman’s ⍴ = -0.41, p = 0.005) (Figure 6). The negative correlation is expected, because we expect higher evolutionary rates to correspond to codon pairs with similar tRNA abundances, and thus lower abundance ratio.

**Figure 6.**
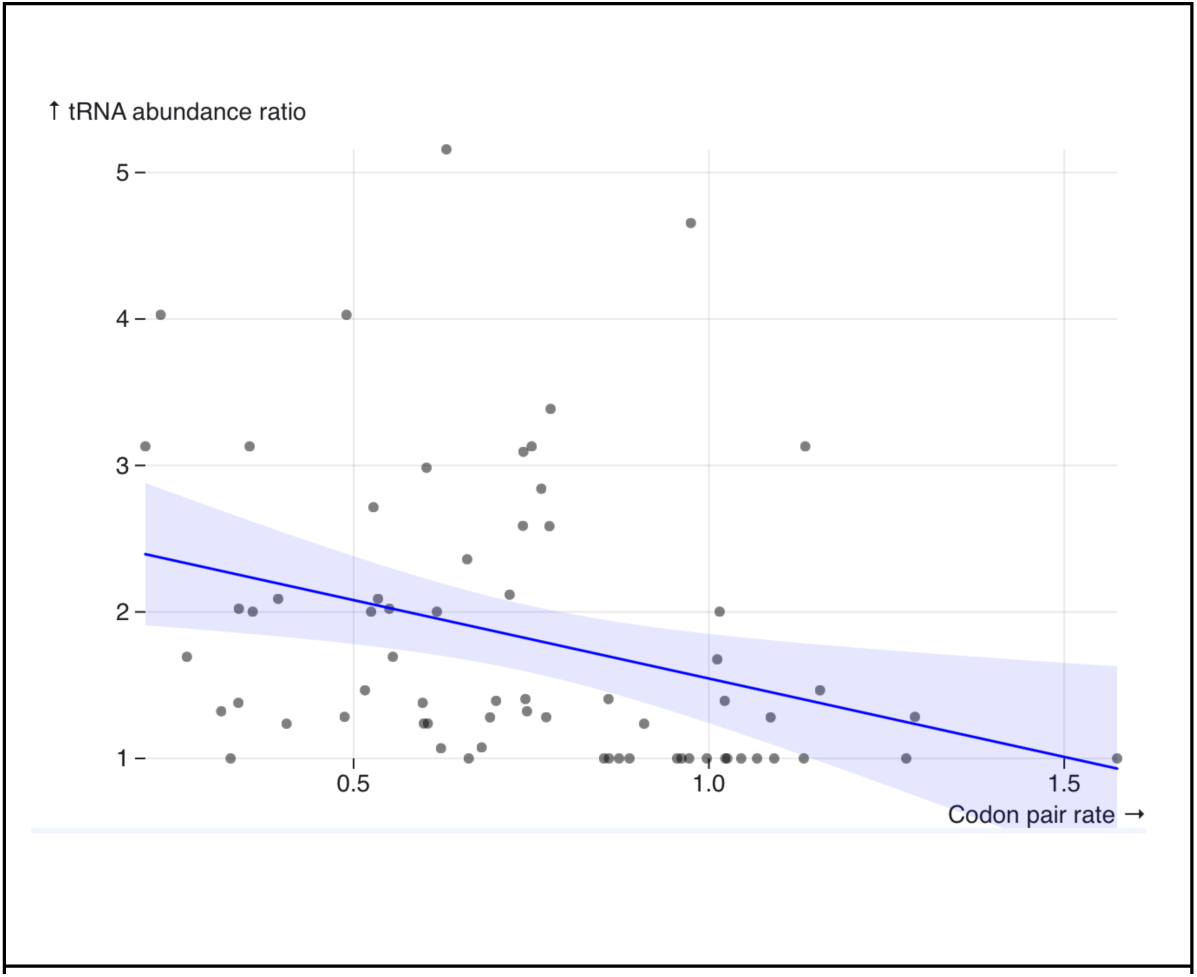
There is a negative correlation (rank-based Spearman’s ρ = −0.41, p = 0.005) between the median inferred synonymous substitution rates (SynREVCodon), and the tRNA abundance ratio for the corresponding codon pair.

### Estimates of dN/dS (⍵) ratios are decreased under MSS models

As demonstrated by Rahman *et al* (Rahman et al. 2021) for simpler MSS models with *a priori* amino-acid partitioning, the estimates of dN/dS ratios are expected to decrease when selection on synonymous substitutions is included in the model. For simpler models, where a subset of codons is designated as neutral and only those substitutions contribute towards dS, the intuition is simple: neutral substitutions occur at faster rates than selected substitutions, and including only those in dS will increase the value of the denominator and thus lower the ratio. Because SynREV and SynREVCodon models use a different parameterization, with dN/dS defined relative to multiple synonymous rates, it is not immediately clear what effect this would have on dN/dS estimates. We compared ⍵ (dN/dS) estimates obtained under the standard MG94xREV model, the SynREV model, and the SynREVCodon model (Figure 7), and found that there is a highly significant (p<<0.001) linear correlation between the estimates obtained under the standard MG94xREV model, and the two MSS models. In both cases, there was a mean reduction ⍵ estimates, by a factor of 0.86 (SynREV) and 0.68 (SynREVCodon).

**Figure 7.**
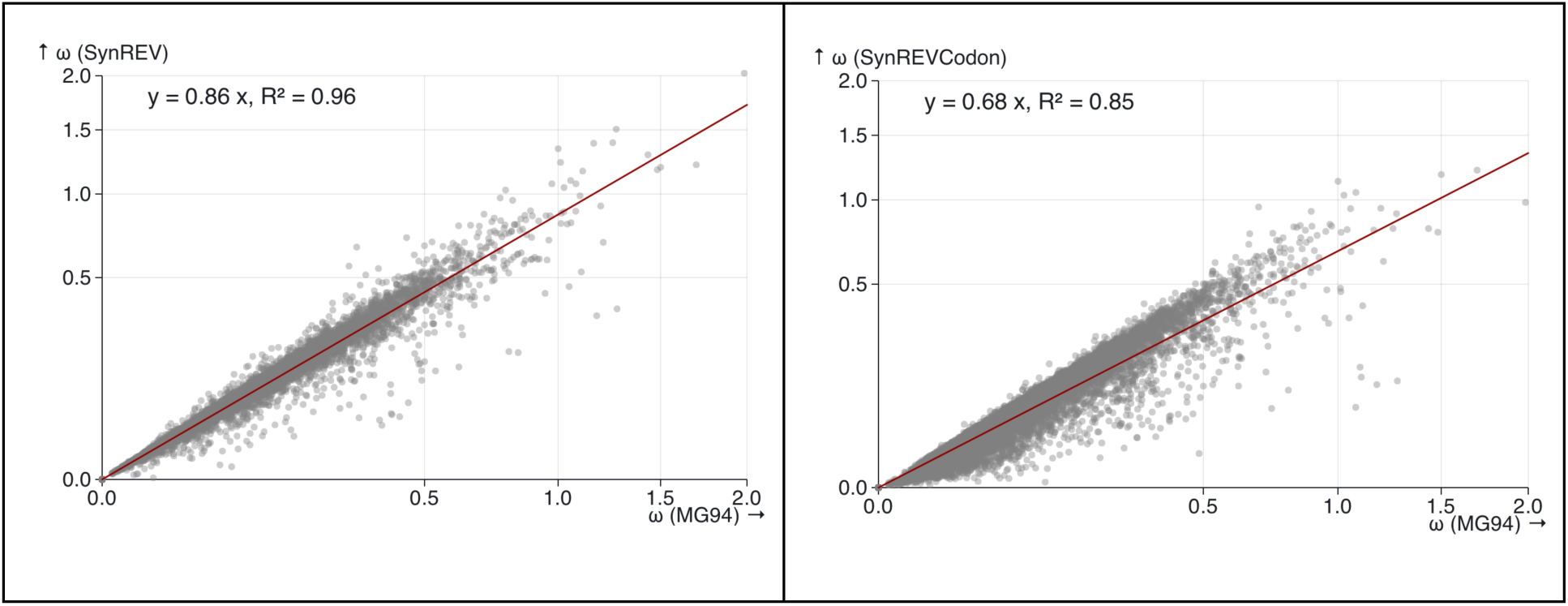
The relationship between ⍵ (dN/dS) under MG94xREV, SynREV, and SynREVCodon models. Linear regression with the 0 intercept was run using the lm (y∼x+0) function in R,

We also compared ⍵ (dN/dS) estimates obtained under the standard MG94xREV model, the SynREV model, and the SynREVCodon model for a set of 14,371 primate gene alignments, containing a median of 25 (range 8-26) sequences with a median of 466 (range 44-5137) codons, and median tree length of 0.40 (range 0.045-1.9) substitutions per nucleotide site. We found that ⍵ estimates were reduced by a factor of 0.895 under the SynREV model, and by a factor of 0.82 under the SynREVCodon model (Figure 8), suggesting that selection on synonymous codon usage is present across taxa. Selection on synonymous codon usage in primates has been reported in the past, even though it may be less pronounced than in some other clades (Chamary et al. 2006; Drummond and Wilke 2008; Dhindsa et al. 2020). As discussed in Rahman *et al* (Rahman et al. 2021), potentially biased dN/dS estimates with similar extent of bias will affect many downstream analyses which rely on them.

**Figure 8.**
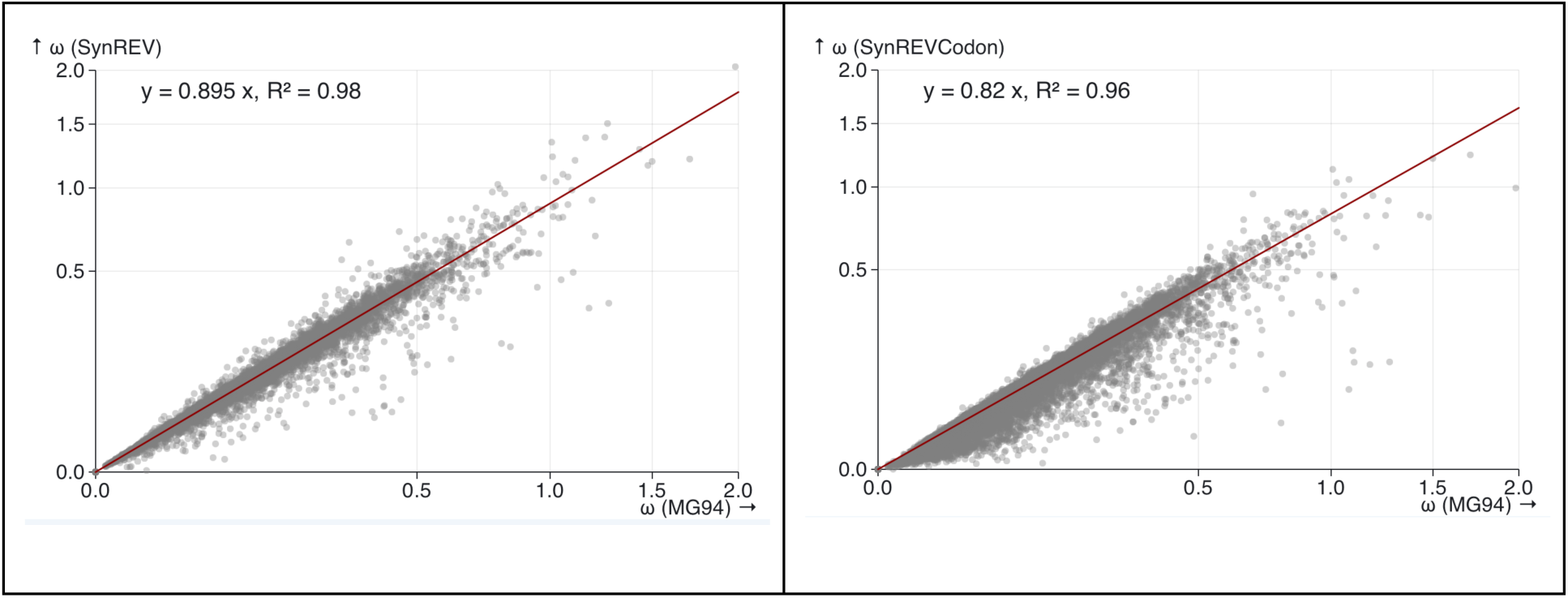
The relationship between ⍵ under MG94xREV, SynREV, and SynREVCodon models for primate gene alignments. Linear regression with the 0 intercept was run using the lm(y∼x+0) function in R.

### Joint estimation of MSS models

Because genes are relatively short, and the number of sequences is small, we expect these rate estimates to be noisy. Indeed, as can be seen in Figure 2, for many genes individual codon pair rate estimates lie in the extreme tails of the distribution: exactly 0 (which is not biologically plausible: surely any codon pair can be exchanged at some, albeit low, rate), or a large value (possible saturation). If we assume that a single MSS model can adequately represent substitution patterns in a large fraction of the genes, then we could simply infer such a model from a large number of genes jointly. This is done while allowing each alignment to use a separate set of parameters such as base frequencies, nucleotide substitution biases, dN/dS ratios, *etc*. Of course, if no such single model exists, *i.e.*, sets of genes have discordant substitution preferences, then a single “average” model may be a poor fit to any individual gene.

From all N=11,844 available alignments, we selected random subsets of K=4, 8, 16, 32, 64, 128, 256, 512, 1024, and 2048 genes (5 independent draws for each K), and jointly inferred MSS models from them. This procedure is useful in establishing how stable rate inferences are based on the number of genes, and between replicates, and also in providing a means for statistically uncontaminated testing of model fit. Indeed, we may use a model inferred from a set of K genes, on the independent (hold-out) set of N-K remaining alignments for validation.

Starting with as few as 8 alignments, MSS models offer a statistically significant improvement over baseline (MG94 models) based on LRT comparisons, and with 16 alignments, they also show a consistent improvement in AIC scores. As the number of alignments increase, the improvement in fit (AIC points) per alignment is around 3.7 for SynREV and 10 for SynREVCodon. Thus, using a “global” MSS model on multiple alignments captures the general substitution patterns of a large fraction of genes.

#### A more detailed examination of how model choice impacts dN/dS estimation

Our MG94-based models define dN/dS indirectly (Table 1), by estimating the non-synonymous substitution rate β and the synonymous substitution rate ɑ separately. The ratio of β/ɑ defines ⍵. One can also define dS (dN) as the expected numbers of synonymous (non-synonymous) substitutions per unit time normalized by the expected number of synonymous and non-synonymous sites. The latter two quantities are non-trivial to define and how this is done may affect dS and dN estimates substantially (see Muse 1996). For the standard MG94 model ⍵ = β, since ɑ := 1. For the MSS and GA-based models (Table 2), because one of the ɑ rates is identified as “neutral”, *i.e.*, constrained to be 1, ⍵ will be defined relative to that rate (referred to as dSn in Rahman et al. 2021). For SynREV and SynREVCodon models, only the mean of ɑ is constrained to be 1, and there is no analogous set of neutral synonymous substitutions.

Let us consider how the estimates of ⍵ vary under different models for the *barren* Drosophila gene (FBgn0014127) in Table 7. In addition to the previously defined models, we also define two new models for illustration purposes.

1. *Joint codon empirical*. This model takes the numerical estimates for ɑ rates from the SynREVCodon model inferred jointly from 1024 alignments (not including FBgn0014127), and then treats them as constant. Such a model is similar to protein empirical models (*e.g.*, Dayhoff et al. 1978; Whelan and Goldman 2001) and the ECM model of Kosiol et al. (Kosiol et al. 2007)
2. *Joint codon discretized*. Using the same joint model as above, the estimates for ɑ are ranked, and the top 15% (by numerical value of the estimates) are taken to be “neutral”, thereby generating an a priori model, with a “neutral” class, and the corresponding dSn rate.

**Table 7.** Estimates of ⍵ under different models for the alignment of 10 sequences for the *barren* gene (FBgn0014127). LRT p-value is computed using MG94 as null, with the appropriate number of degrees of freedom for each model (17, 67, 0, 1). Std.dev [ɑ] is computed directly on the distribution of 68 codon-pair rates for each model.

| Model | $\omega$ | $\log(L)$ | LRT p-value | Std.dev [$\alpha$] |
| --- | --- | --- | --- | --- |
| MG94 | 0.085 | -7040.8 | n/a | 0.000 |
| SynREV | 0.074 | -7023.43 | 0.007 | 0.268 |
| SynREVCodon | 0.063 | -6982.27 | 0.0002 | 0.744 |
| Joint codon empirical | 0.085 | -7026.40 | $<10^{-6}$ | 0.176 |
| Joint codon discretized | 0.041 | -7024.58 | $<10^{-6}$ | 0.200 |

Both of these models can be viewed as practical alternatives to the rather costly and noisy gene-by-gene or joint inference. Because these models are precomputed, they can be applied to genes from the same genus to obtain a first-order correction for variable ɑ rates with a relatively minimal parametric or computational overhead.

The estimate of ⍵ is sensitive to the choice of model, and is no greater than the MG94 (baseline) estimate for any of the MSS models. Biological datasets appear to support a significant but not extreme reduction (0.6-0.8x) in ⍵ estimates compared to the MG94 model.

To establish a statistical performance baseline we ran an ⍵ bias simulation experiment. We randomly selected 100 *Drosophila* alignments, and then used the MG94, SynREV, and SynREVCodon model fits for each of these alignments to simulate 100 replicates per model (300 total per alignment). We then fitted MG94, SynREV, and SynREVCodon to each of the 30,000 replicates to estimate ⍵ and assess the degree of ⍵ estimation bias (Table 8).

1. Estimates obtained under the MG94 model are unbiased when the generating model is also MG94, and have a strong upward bias when the generating model is an MSS model. This bias is often severe.
2. Estimates obtained under the SynREVCodon model are biased downwards even when the generating model is also SynREVCodon; they are quite noisy (high variance), as can be expected for such a parameter rich model. Relative bias is less extreme than the corresponding values for the MG94 model.
3. The behavior of estimates obtained under the SynREV model is somewhere in between; a small downward bias when the true model is MG94, a significant upward bias but less extreme than MG94 on the data from SynREVCodon, and a slight downward bias on the data generated under the SynREV model itself.

**Table 8.** Relative bias, defined as (*E*[ω̂] − ω_*true*_)/ω_*true*_), for different generating and fitted model pairs. Rate of severe bias is defined as the proportion of simulation scenarios where the true value of ⍵ was outside the [2.5% - 97.5%] quantile distribution for ⍵ estimates, i.e., the true value is in the tails of the sampling distribution.

| Generating Mode | Fitted Model | $\omega$ estimate relative bias | | |
| --- | --- | --- | --- | --- |
|  |  | Mean | Variance | Rate of severe* bias |
| MG94 | MG94 | 0.001 | 0.015 | 0.00% |
| MG94 | SYNREV | -0.022 | 0.049 | 0.00% |
| MG94 | SYNREVCODON | -0.135 | 0.126 | 0.00% |
| SYNREV | MG94 | 0.175 | 0.575 | 9.09% |
| SYNREV | SYNREV | -0.023 | 0.058 | 0.00% |
| SYNREV | SYNREVCODON | -0.049 | 0.187 | 2.02% |
| SYNREVCODON | MG94 | 0.665 | 1.225 | 61.62% |
| SYNREVCODON | SYNREV | 0.351 | 0.575 | 25.25% |
| SYNREVCODON | SYNREVCODON | -0.107 | 0.137 | 0.00% |

Fitting complex models to small alignments introduces statistical inefficiencies and biases, even though the overall pattern is clear. MG94 severely overestimates dN/dS when the true model is an MSS model, and although both MSS models show downward estimator bias, it is much less extreme than the MG94 upward bias.

What is clear, however, is that even when estimated as an alignment-wide parameter, ⍵, is rather sensitive to how one chooses to model the variation in ɑ rates. While we do not examine it further here, previous work in Rahman et al. 2021 shows that ⍵ estimates are even more sensitive in models which allow ⍵ to vary across sites and branches under the *a priori* MSS models.

## Discussion

We developed a broad class of phylogenetic codon-substitution models to infer relative synonymous substitution rates from coding sequence alignments. These multiclass synonymous substitution (MSS) models expand upon our prior work (Rahman et al. 2021), by inferring substitution rate classes directly from the data, instead of relying on predefined classes of “neutral” or “selected” substitutions. When applied to gene alignments from twelve closely related *Drosophila* species, the MSS models reveal a striking degree of heterogeneity in synonymous substitution rates across the genes. MSS models provide a significantly better fit to the data compared to standard models where all synonymous substitutions share the same rate. This is true even for the fine-grained SynREVCodon model that endows individual synonymous codon pairs with separate rates, an indicator that the standard modeling assumptions describe the data quite poorly and there is ample room for improvement. Goodness-of-fit improvements over standard substitution models are easy to obtain and speak more to the gross inadequacy of existing models than to the biological veracity of the newer ones (Delport, Scheffler, Gravenor, et al. 2010; Jones et al. 2018). We corroborated the inferences made by the MSS models using forward population genetic simulations (an entirely different modeling framework), using an external dataset of within-population SNP variation (an independent source of a different type of variation data, with within-species and between-species correlation expected), and by confirming that the ranking of inferred rates is in accordance with relative tRNA abundance (a key mechanistic factor).

MSS models reveal a spectrum of substitution rates between synonymous codons across *Drosophila* genes, with a significant (several-fold) difference between the slowest and the fastest rates. The inferred distribution of rates shows, as expected, that substitutions between codons whose corresponding tRNAs occur at significantly different abundances tend to have lower rates, likely attributable to selection for translational efficiency. However, many other factors, both genome-wide, and gene-level likely contribute to this variation. Further elucidation of these factors requires substantial additional work, and likely relies on data that are not readily available.

MSS models, like all other models, are approximations of the true biological process and are misspecified. For example, MSS models do not account for time-variable synonymous substitution rates, which can confound interpretations of model fit in mutation-selection models (Jones et al. 2018). Additionally, the interaction of context-dependent substitution processes and emergent non-Markovian dynamics presents further challenges that are not explicitly addressed in our model. While these factors are undoubtedly important, incorporating them into a comprehensive framework would significantly increase the complexity of the model, and lead to loss of computational feasibility and statistical performance. Accounting for additional biological phenomena would undoubtedly improve the model’s fit to the data and the overall realism of the model, but the feasibility of such an endeavor is the subject of future work. Nevertheless, we believe that the MSS modeling framework provides a useful and practical tool for capturing the key drivers of synonymous substitution and offers valuable insights into the evolutionary processes shaping codon usage.

The removal of a long-held modeling assumption regarding the constancy of synonymous substitution rates across codon pairs will undoubtedly influence inferences of other evolutionary parameters. We showed that the introduction of multiclass synonymous substitutions reduces the estimates of gene-wide dN/dS, similar to what was reported in Rahman *et al* (Rahman et al. 2021) for simpler *a priori* models, thereby affecting downstream analyses which rely on dN/dS or its derivatives. Additional analyses will be required to evaluate the impact that MSS models may have on other common tasks, such as phylogenetic inference, or time tree estimation. Our work here contributes to the recent thread in evolutionary modeling literature which re-examines historically accepted assumptions (Wisotsky et al. 2020; Lucaci et al. 2021), and the influence of model choice on different types of analyses which require the selection of a substitution model.

We envision that the MSS framework presented here will have both short-term and longer-term influence on comparative evolutionary analyses. In the short term, MSS models can be applied to genomes from different organisms and clades to infer and catalog the distributions of substitution rates among synonymous codons. Identifying a robust subset of synonymous substitutions which evolve in a neutral fashion directly from the data will improve inference that requires a neutral reference, and permit the examination of what factors inform the composition of the neutral set and partition the rates into different evolutionary classes. We also anticipate that a single MSS model will not describe all the genes equally well. The MSS framework can be used to select the genes which deviate from genome-wide averages and then investigate the attributes of these outlier genes. In the longer term, natural extensions to the MSS framework, and incorporation of other processes known to be important can give rise to more biologically informative tools. Specifically, MSS models can be combined with models which allow non-synonymous rates to depend on which codons are being exchanged (Wong et al. 2006). With the exception of a few models (Kosiol et al. 2007; Yang and Nielsen 2008), codon-pair level variation in synonymous and non-synonymous rates have not been studied jointly, and they are likely entangled, and correlated in non-trivial ways. MSS models can be extended to allow different types of site-to-site and branch-to-branch variation to accommodate heterotachy and other complex processes.

Our work contributes to the growing collection of studies revealing that synonymous substitution rates are neither fully neutral, nor accrue at constant rates over sites and branches (P M Sharp and Li 1987; Shields et al. 1988; Iriarte et al. 2013; Trotta 2013; Wisotsky et al. 2020). The genome-wide selective force that powers the MSS models is that of weak selection, and though its effects are small when compared to the effects of non-synonymous substitutions (Rahman et al. 2021), it is nonetheless readily detectable and informative. We hope the collection of tools and motivating examples provided here will entice additional research into the scope, drivers, and effect of non-neutral processes affecting synonymous mutations and substitutions.

### Data Availability

The MSS model is included with the full set of Hyphy analyses, which can be downloaded at https://github.com/veg/hyphy-analyses/tree/master. A SynREVCodon analysis of a single gene can be run as follows:

The joint model fitter script is included in the base Hyphy package. The script requires a text list of newline-separated file paths, where each file contains a codon-aware alignment and the corresponding gene tree. It is run as follows:

The MSS-selector-2 package is also included in the base Hyphy package. Like the joint model fitter script, MSS-selector-2 requires a text list of newline-separated file paths, where each file contains a codon-aware alignment and the corresponding gene tree. It also requires that users specify the number of classes into which codons shall be partitioned. The script is run as follows:

Code to run forward simulations can be found at https://github.com/ampivirotto/mss_forward_sim. Simulated alignments used in the forward simulation analysis can be found at https://github.com/hverdonk/MSS-supplemental/tree/main/forward_simulations, and the simulated population tree is in Figure S1. Scripts and data used in all other analyses can also be found at https://github.com/hverdonk/MSS-supplemental/.

## Supporting information

Genetic algorithm model fits on parametric simulation data

Drosophila genome reference IDs and access date

Drosophila codon usage patterns

## Acknowledgements

We would like to thank Avery Selberg, Steven Weaver, Stephen Shank, and all other members of the ACME lab for their insightful feedback and discussions.

## Bibliography

1. Akashi H. 1994. Synonymous codon usage in Drosophila melanogaster: natural selection and translational accuracy. Genetics 136:927–935.

2. Armstrong J, Hickey G, Diekhans M, Fiddes IT, Novak AM, Deran A, Fang Q, Xie D, Feng S, Stiller J, et al. 2020. Progressive Cactus is a multiple-genome aligner for the thousand-genome era. Nature 587:246–251.

3. Behrens A, Rodschinka G, Nedialkova DD. 2021. High-resolution quantitative profiling of tRNA abundance and modification status in eukaryotes by mim-tRNAseq. Mol. Cell 81:1802–1815.e7.

4. Chamary JV, Parmley JL, Hurst LD. 2006. Hearing silence: non-neutral evolution at synonymous sites in mammals. Nat. Rev. Genet. 7:98–108.

5. Chiari Y, Dion K, Colborn J, Parmakelis A, Powell JR. 2010. On the Possible Role of tRNA Base Modifications in the Evolution of Codon Usage: Queuosine and Drosophila. J. Mol. Evol. 70:339–345.

6. Crick FHC. 1966. Codon—anticodon pairing: The wobble hypothesis. J. Mol. Biol. 19:548–555.

7. Dayhoff MO, Schwartz RM, Orcutt BC. 1978. A model of evolutionary change in proteins. In: Atlas of protein sequence and structure. Vol. 5. Washington, D.C.: Biomedical Research Foundation. p. 345–352.

8. Delport W, Scheffler K, Botha G, Gravenor MB, Muse SV, Pond SLK. 2010. CodonTest: Modeling Amino Acid Substitution Preferences in Coding Sequences. PLOS Comput. Biol. 6:e1000885.

9. Delport W, Scheffler K, Gravenor MB, Muse SV, Pond SK. 2010. Benchmarking Multi-Rate Codon Models. PLOS ONE 5:e11587.

10. Dhindsa RS, Copeland BR, Mustoe AM, Goldstein DB. 2020. Natural Selection Shapes Codon Usage in the Human Genome. Am. J. Hum. Genet. 107:83–95.

11. Drummond DA, Wilke CO. 2008. Mistranslation-Induced Protein Misfolding as a Dominant Constraint on Coding-Sequence Evolution. Cell 134:341–352.

12. Eshelman LJ. 1991. The CHC Adaptive Search Algorithm: How to Have Safe Search When Engaging in Nontraditional Genetic Recombination. In: Rawlins GJE, editor. Foundations of Genetic Algorithms. Vol. 1. Elsevier. p. 265–283. Available from: https://www.sciencedirect.com/science/article/pii/B9780080506845500203

13. Ewens WJ. 2004. Mathematical population genetics [electronic resource] : 1. Theoretical introduction. New York : Springer Available from: http://archive.org/details/springer_10.1007-978-0-387-21822-9

14. Felsenstein J. 1981. Evolutionary trees from DNA sequences: A maximum likelihood approach. J. Mol. Evol. 17:368–376.

15. Goetz RM, Fuglsang A. 2005. Correlation of codon bias measures with mRNA levels: analysis of transcriptome data from Escherichia coli. Biochem. Biophys. Res. Commun. 327:4–7.

16. Hey J, Kliman RM. 2002. Interactions between natural selection, recombination and gene density in the genes of Drosophila. Genetics 160:595–608.

17. Ikemura T. 1981. Correlation between the abundance of Escherichia coli transfer RNAs and the occurrence of the respective codons in its protein genes: A proposal for a synonymous codon choice that is optimal for the E. coli translational system. J. Mol. Biol. 151:389–409.

18. Iriarte A, Baraibar JD, Romero H, Castro-Sowinski S, Musto H. 2013. Evolution of optimal codon choices in the family Enterobacteriaceae. Microbiology 159:555–564.

19. Jones CT, Youssef N, Susko E, Bielawski JP. 2018. Phenomenological Load on Model Parameters Can Lead to False Biological Conclusions. Mol. Biol. Evol. 35:1473–1488.

20. Kanaya S, Yamada Y, Kinouchi M, Kudo Y, Ikemura T. 2001. Codon Usage and tRNA Genes in Eukaryotes: Correlation of Codon Usage Diversity with Translation Efficiency and with CG-Dinucleotide Usage as Assessed by Multivariate Analysis. J. Mol. Evol. 53:290–298.

21. Kimura M. 1962. ON THE PROBABILITY OF FIXATION OF MUTANT GENES IN A POPULATION. Genetics 47:713–719.

22. Kosakovsky Pond SL, Frost SDW. 2005. Not So Different After All: A Comparison of Methods for Detecting Amino Acid Sites Under Selection. Mol. Biol. Evol. 22:1208–1222.

23. Kosakovsky Pond SL, Mannino FV, Gravenor MB, Muse SV, Frost SDW. 2007. Evolutionary Model Selection with a Genetic Algorithm: A Case Study Using Stem RNA. Mol. Biol. Evol. 24:159–170.

24. Kosakovsky Pond SL, Posada D, Gravenor MB, Woelk CH, Frost Simon D. W. 2006. Automated Phylogenetic Detection of Recombination Using a Genetic Algorithm. Mol. Biol. Evol. 23:1891–1901.

25. Kosakovsky Pond SL, Posada D, Gravenor MB, Woelk CH, Frost Simon D.W. 2006. GARD: a genetic algorithm for recombination detection. Bioinformatics 22:3096–3098.

26. Kosiol C, Holmes I, Goldman N. 2007. An Empirical Codon Model for Protein Sequence Evolution. Mol. Biol. Evol. 24:1464–1479.

27. Lack JB, Cardeno CM, Crepeau MW, Taylor W, Corbett-Detig RB, Stevens KA, Langley CH, Pool JE. 2015. The Drosophila Genome Nexus: A Population Genomic Resource of 623 Drosophila melanogaster Genomes, Including 197 from a Single Ancestral Range Population. Genetics 199:1229–1241.

28. Lucaci AG, Wisotsky SR, Shank SD, Weaver S, Pond SLK. 2021. Extra base hits: Widespread empirical support for instantaneous multiple-nucleotide changes. PLOS ONE 16:e0248337.

29. Marais G, Domazet-Lošo T, Tautz D, Charlesworth B. 2004. Correlated Evolution of Synonymous and Nonsynonymous Sites in Drosophila. J. Mol. Evol. 59:771–779.

30. Marais G, Duret L. 2001. Synonymous Codon Usage, Accuracy of Translation, and Gene Length in Caenorhabditis elegans. J. Mol. Evol. 52:275–280.

31. Moriyama EN, Powell JR. 1997. Codon Usage Bias and tRNA Abundance in Drosophila. J. Mol. Evol. 45:514–523.

32. Muse SV. 1996. Estimating synonymous and nonsynonymous substitution rates. Mol. Biol. Evol. 13:105–114.

33. Muse SV, Gaut BS. 1994. A likelihood approach for comparing synonymous and nonsynonymous nucleotide substitution rates, with application to the chloroplast genome. Mol. Biol. Evol. 11:715–724.

34. Pivirotto A. 2024. ampivirotto/mss_forward_sim. Available from: https://github.com/ampivirotto/mss_forward_sim

35. Pond SK, Delport W, Muse SV, Scheffler K. 2010. Correcting the Bias of Empirical Frequency Parameter Estimators in Codon Models. PLOS ONE 5:e11230.

36. Rahman S, Kosakovsky Pond SL, Webb A, Hey J. 2021. Weak selection on synonymous codons substantially inflates dN/dS estimates in bacteria. Proc. Natl. Acad. Sci. 118:e2023575118.

37. Sharp P M, Li WH. 1987. The rate of synonymous substitution in enterobacterial genes is inversely related to codon usage bias. Mol. Biol. Evol. 4:222–230.

38. Sharp Paul M., Li W-H. 1987. The codon adaptation index-a measure of directional synonymous codon usage bias, and its potential applications. Nucleic Acids Res. 15:1281–1295.

39. Sharp SJ, Schaack J, Cooley L, Burke DJ, Soil D. 1985. Structure and Transcription of Eukaryotic tRNA Gene. Crit. Rev. Biochem. 19:107–144.

40. Shields DC, Sharp PM, Higgins DG, Wright F. 1988. “Silent” sites in Drosophila genes are not neutral: evidence of selection among synonymous codons. Mol. Biol. Evol. 5:704–716.

41. Suvorov A, Kim BY, Wang J, Armstrong EE, Peede D, D’Agostino ERR, Price DK, Waddell PJ, Lang M, Courtier-Orgogozo V, et al. 2022. Widespread introgression across a phylogeny of 155 *Drosophila* genomes. Curr. Biol. 32:111–123.e5.

42. Torres AG, Piñeyro D, Rodríguez-Escribà M, Camacho N, Reina O, Saint-Léger A, Filonava L, Batlle E, Ribas de Pouplana L. 2015. Inosine modifications in human tRNAs are incorporated at the precursor tRNA level. Nucleic Acids Res. 43:5145–5157.

43. Trotta E. 2013. Selection on codon bias in yeast: a transcriptional hypothesis. Nucleic Acids Res. 41:9382–9395.

44. Whelan S, Goldman N. 2001. A General Empirical Model of Protein Evolution Derived from Multiple Protein Families Using a Maximum-Likelihood Approach. Mol. Biol. Evol. 18:691–699.

45. Wisotsky SR, Kosakovsky Pond SL, Shank SD, Muse SV. 2020. Synonymous Site-to-Site Substitution Rate Variation Dramatically Inflates False Positive Rates of Selection Analyses: Ignore at Your Own Peril. Mol. Biol. Evol. 37:2430–2439.

46. Wong WS, Sainudiin R, Nielsen R. 2006. Identification of physicochemical selective pressure on protein encoding nucleotide sequences. BMC Bioinformatics 7:148.

47. Yang Z, Nielsen R. 2008. Mutation-Selection Models of Codon Substitution and Their Use to Estimate Selective Strengths on Codon Usage. Mol. Biol. Evol. 25:568–579.

48. Zhang W, Foo M, Eren AM, Pan T. 2022. tRNA modification dynamics from individual organisms to metaepitranscriptomics of microbiomes. Mol. Cell 82:891–906.

