## Supplemental Figures and Tables for "A new comparative framework for estimating selection on synonymous substitutions"

### Supplement

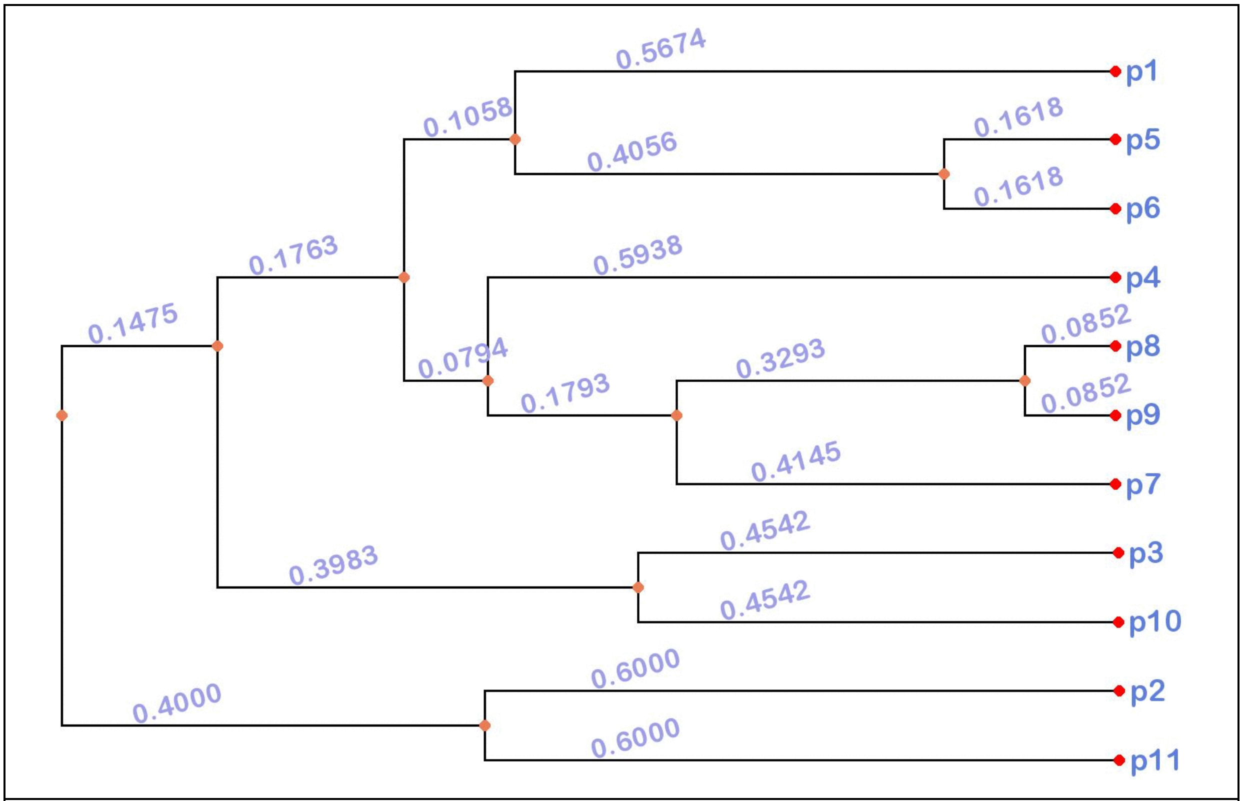


Figure S1. Phylogeny of population splitting events used by the forward simulator

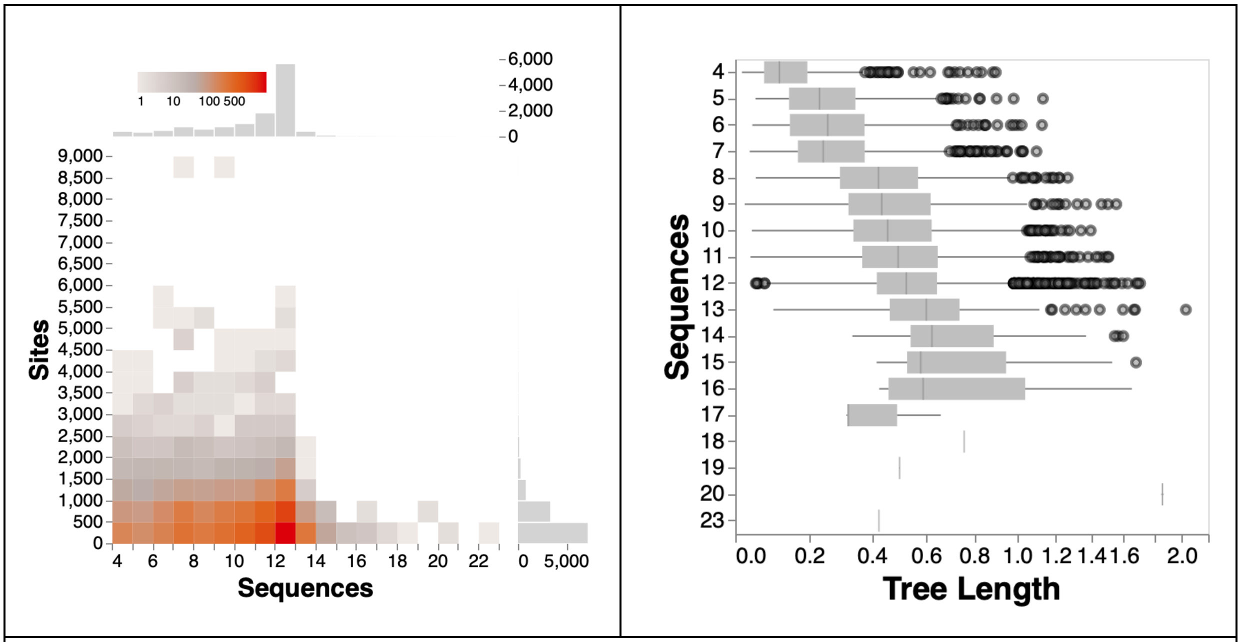


Figure S2. (Left) Dimensions of individual alignments. (Right) Tree lengths (expected substitutions / site) by the number of sequences.

| Amino Acid | Neutral Set(s) |
|---|---|
| A | GCG GCT GCA GCC |
| C | TGT |
| C | TGC |
| D | GAC |
| D | GAT |
| E | GAA GAG |
| F | TTT TTC |
| G | GGA GGC GGT |
| G | GGG |
| H | CAC |
| H | CAT |
| I | ATT |
| I | ATC ATA |
| K | AAA |
| K | AAG |
| L | CTT CTA CTG CTC |
| L | TTG TTA |
| N | AAC |
| N | AAT |
| P | CCA CCC CCT CCG |
| Q | CAA |
| Q | CAG |
| R | CGA AGA AGG CGG |
| R | CGC CGT |

| | |
|---|---|
| S | AGT TCG TCT AGC TCA TCC |
| T | ACC ACT |
| T | ACG ACA |
| V | GTT GTC |
| V | GTA GTG |
| Y | TAC TAT |

**Table S1:** True model for forward simulations. Each amino acid may have one or two classes of neutral synonymous substitutions, with each class' member codons represented by a separate row in the table. Substitutions within a class (row) are treated as neutral, synonymous substitutions between classes (rows) are treated as selected.

| **Amino acid** | **Codon 1** | **Codon 2** | **Total subs** | **Per alignment** | | | **Alignments with non-zero counts** |
|---|---|---|---|---|---|---|---|
| | | | | ***Mean*** | ***Median*** | ***Maximum*** | |
| **A** | GCA | GCC | 59754 | 5.05 | 3 | 86 | 10328 |
| **A** | GCA | GCG | 58383 | 4.93 | 3 | 103 | 10030 |
| **A** | GCA | GCT | 34207 | 2.89 | 2 | 58 | 8729 |
| **A** | GCC | GCG | 52474 | 4.43 | 3 | 67 | 9822 |
| **A** | GCC | GCT | 114063 | 9.63 | 7 | 174 | 11044 |
| **A** | GCG | GCT | 30633 | 2.59 | 2 | 59 | 8627 |
| **C** | TGC | TGT | 55691 | 4.70 | 3 | 214 | 9450 |
| **D** | GAC | GAT | 160809 | 13.58 | 10 | 271 | 11257 |
| **E** | GAA | GAG | 178073 | 15.03 | 10 | 352 | 11278 |
| **F** | TTC | TTT | 151186 | 12.76 | 9 | 255 | 11165 |
| **G** | GGA | GGC | 79074 | 6.68 | 5 | 126 | 10598 |
| **G** | GGA | GGG | 46943 | 3.96 | 3 | 80 | 9618 |
| **G** | GGA | GGT | 48845 | 4.12 | 3 | 135 | 9597 |
| **G** | GGC | GGG | 30676 | 2.59 | 2 | 36 | 8779 |
| **G** | GGC | GGT | 107251 | 9.06 | 6 | 155 | 10931 |
| **G** | GGG | GGT | 17310 | 1.46 | 1 | 27 | 7083 |
| **H** | CAC | CAT | 78022 | 6.59 | 4 | 99 | 10381 |
| **I** | ATA | ATC | 41644 | 3.52 | 2 | 66 | 9319 |
| **I** | ATA | ATT | 43272 | 3.65 | 2 | 141 | 9057 |

| | | | | | | | |
|---|---|---|---|---|---|---|---|
| **I** | ATC | ATT | 120581 | 10.18 | 7 | 180 | 11079 |
| **K** | AAA | AAG | 150628 | 12.72 | 9 | 229 | 11198 |
| **L** | CTA | CTC | 25321 | 2.14 | 1 | 58 | 8102 |
| **L** | CTA | CTG | 104567 | 8.83 | 6 | 185 | 10822 |
| **L** | CTA | CTT | 24548 | 2.07 | 1 | 72 | 7431 |
| **L** | CTA | TTA | 24248 | 2.05 | 1 | 83 | 7173 |
| **L** | CTC | CTG | 85905 | 7.25 | 5 | 153 | 10515 |
| **L** | CTC | CTT | 73776 | 6.23 | 4 | 159 | 10470 |
| **L** | CTG | CTT | 59018 | 4.98 | 3 | 122 | 9945 |
| **L** | CTG | TTG | 159346 | 13.45 | 9 | 281 | 11225 |
| **L** | TTA | TTG | 39966 | 3.37 | 2 | 134 | 8460 |
| **N** | AAC | AAT | 137345 | 11.60 | 8 | 284 | 11159 |
| **P** | CCA | CCC | 48959 | 4.13 | 3 | 59 | 9848 |
| **P** | CCA | CCG | 77699 | 6.56 | 4 | 101 | 10398 |
| **P** | CCA | CCT | 25403 | 2.14 | 1 | 52 | 7612 |
| **P** | CCC | CCG | 51863 | 4.38 | 3 | 55 | 9832 |
| **P** | CCC | CCT | 59841 | 5.05 | 3 | 78 | 10073 |
| **P** | CCG | CCT | 25826 | 2.18 | 1 | 38 | 7930 |
| **Q** | CAA | CAG | 128038 | 10.81 | 7 | 247 | 10992 |
| **R** | AGA | AGG | 36025 | 3.04 | 2 | 73 | 8492 |
| **R** | AGA | CGA | 18976 | 1.60 | 1 | 40 | 6868 |
| **R** | AGG | CGG | 30640 | 2.59 | 2 | 55 | 8405 |
| **R** | CGA | CGC | 38502 | 3.25 | 2 | 58 | 9190 |
| **R** | CGA | CGG | 51359 | 4.34 | 3 | 68 | 9558 |
| **R** | CGA | CGT | 25308 | 2.14 | 1 | 43 | 7866 |
| **R** | CGC | CGG | 41889 | 3.54 | 2 | 49 | 9184 |
| **R** | CGC | CGT | 72197 | 6.10 | 4 | 109 | 10286 |
| **R** | CGG | CGT | 22725 | 1.92 | 1 | 26 | 7752 |
| **S** | AGC | AGT | 81440 | 6.88 | 4 | 143 | 10417 |
| **S** | TCA | TCC | 31573 | 2.67 | 2 | 57 | 8789 |
| **S** | TCA | TCG | 52827 | 4.46 | 3 | 107 | 9750 |
| **S** | TCA | TCT | 14331 | 1.21 | 0 | 44 | 5808 |
| **S** | TCC | TCG | 57298 | 4.84 | 3 | 82 | 9984 |
| **S** | TCC | TCT | 61194 | 5.17 | 3 | 125 | 10179 |
| **S** | TCG | TCT | 24585 | 2.08 | 1 | 51 | 7847 |
| **T** | ACA | ACC | 43368 | 3.66 | 2 | 63 | 9583 |

| | | | | | | | |
|---|---|---|---|---|---|---|---|
| **T** | ACA | ACG | 56548 | 4.77 | 3 | 84 | 9846 |
| **T** | ACA | ACT | 26158 | 2.21 | 1 | 64 | 7835 |
| **T** | ACC | ACG | 44578 | 3.76 | 2 | 65 | 9450 |
| **T** | ACC | ACT | 70086 | 5.92 | 4 | 109 | 10513 |
| **T** | ACG | ACT | 27903 | 2.36 | 1 | 55 | 8206 |
| **V** | GTA | GTC | 18135 | 1.53 | 1 | 34 | 7295 |
| **V** | GTA | GTG | 68488 | 5.78 | 4 | 128 | 10356 |
| **V** | GTA | GTT | 20096 | 1.70 | 1 | 80 | 7120 |
| **V** | GTC | GTG | 60431 | 5.10 | 4 | 78 | 10160 |
| **V** | GTC | GTT | 58191 | 4.91 | 3 | 107 | 10320 |
| **V** | GTG | GTT | 56655 | 4.78 | 3 | 94 | 10129 |
| **Y** | TAC | TAT | 99082 | 8.37 | 6 | 168 | 10865 |

**Table S2**. Descriptive statistics for each of the 67 SNSS substitutions in the *Drosophila* dataset, sorted by the encoded amino-acid.

#### Genetic algorithm testing: parametric simulations under the MG94 (null) model

The genetic algorithm performance was tested on null data through simulation of 1500 alignments under the Muse-Gaut 1994 (MG94) model. We generated simulated alignments under the empirical tree topology for the *Enterobacteriaceae* atpD gene alignment, which contains 15 taxa. Subsequently, we ran the genetic algorithm on 25 bootstrap replicates, each consisting of 100 alignments. The selection of the “best” model, either the null model or any alternate models, was evaluated for each replicate using the Bayesian Information Criterion (BIC) calculated for each model during the genetic algorithm procedure. BIC scores for each replicate can be found in `GA_results_for_MG94_data.csv`.
